# Cell cycle dynamics of chromosomal organisation at single-cell resolution

**DOI:** 10.1101/094466

**Authors:** Takashi Nagano, Yaniv Lubling, Csilla Varnai, Carmel Dudley, Wing Leung, Yael Baran, Netta Mandelson-Cohen, Steven Wingett, Peter Fraser, Amos Tanay

## Abstract

Chromosomes in proliferating metazoan cells undergo dramatic structural metamorphoses every cell cycle, alternating between a highly condensed mitotic structure facilitating chromosome segregation, and a decondensed interphase structure accommodating transcription, gene silencing and DNA replication. These cyclical structural transformations have been evident under the microscope for over a century, but their molecular-level analysis is still lacking. Here we use single-cell Hi-C to study chromosome conformations in thousands of individual cells, and discover a continuum of *cis*-interaction profiles that finely position individual cells along the cell cycle. We show that chromosomal compartments, topological domains (TADs), contact insulation and long-range loops, all defined by ensemble Hi-C maps, are governed by distinct cell cycle dynamics. In particular, DNA replication correlates with build-up of compartments and reduction in TAD insulation, while loops are generally stable from G1 through S and G2. Analysing whole genome 3D structural models using haploid cell data, we discover a radial architecture of chromosomal compartments with distinct epigenomic signatures. Our single-cell data creates an essential new paradigm for the re-interpretation of chromosome conformation maps through the prism of the cell cycle.

---

As cells progress from mitosis to interphase and back again, their chromosomes undergo dramatic structural alterations that have intrigued microscopists for over a century^1^. In preparation for mitosis and cell division chromosomes are packaged into rod-like structures that are functionally inert. Upon mitotic exit chromosomes rapidly decondense into structures that allow regulated access to functional elements enabling genome functions such as transcription, gene silencing and DNA replication. Though the structural and organizational principles of the interphase genome have been the subject of intense study, genome wide, high-resolution understanding of chromosome cell cycle dynamics is still limited by current technologies.

The development of high-throughput chromosome conformation capture (Hi-C) technologies opened up the possibility of genome-scale mapping of chromosome conformation by massive statistical analysis of chromosomal contacts. Statistical analysis of Hi-C contact matrices have revealed fundamental properties of chromosomes, including a TAD structure, its correspondence with transcriptionally active (A) and inactive (B) nuclear compartments, and the identification of specific cohesin/CTCF loops anchoring some of these TADs^2–10^. These features shape our current understanding of genome structure and function by linking gene regulation and transcriptional output with physical and cell-type specific properties of the chromosomes that embed them.

Previously, we developed single-cell Hi-C to begin to assess the heterogeneity underlying chromosomal conformations and how they contribute to ensemble Hi-C maps. Our initial set of single-cell maps revealed evidence of TAD organization and other key features such as compartmentalisation at single-cell resolution, indicating that these organisations actually take place at single-cell level^11^. On the other hand, the data also indicated substantial variation in *trans*- and *cis*-chromosomal conformations, suggesting that comprehensive characterization of such heterogeneity is needed in order to understand chromosome structure and function. The cell cycle dynamics of chromosome conformation has recently been studied in cell populations by Hi-C and at specific loci by 4C^12,13^. Both have revealed a loss of TADs and nuclear compartments in mitosis, providing the first molecular details of the dramatic structural alterations associated with mitosis. Nevertheless, these population measurements, even when assessing sorted or synchronised cells, provide an average of highly variable cell-to-cell conformations, limiting high-resolution reconstruction of chromosome dynamics.

Here we scale up single-cell Hi-C using flow cytometry sorting, a new library preparation protocol and automation scheme, and apply this methodology to thousands of diploid and haploid mouse embryonic stem cells (ESC). Having thousands of datasets has allowed us to develop a strategy for in-silico cell cycle phasing of single nuclei, and use it to study the cell cycle dynamics of chromosomal structural features. We also present the first whole genome, restraint-based three-dimensional (3D) models from hundreds of individual G1 phase haploid nuclei. Our results reveal a continuum of dynamic chromosomal structural features throughout the cell cycle, creating a new point of reference for interpretation of Hi-C maps.

## RESULTS

### Improved multiplexed single-cell Hi-C

To improve genomic coverage we redesigned and optimized the efficiency of our original single cell Hi-C approach^11^ to allow effective use of frequent 4-cutter restriction enzymes, and a one-step preparation of library templates with Tn5-transposase tagmentation. To increase throughput we used flow cytometry sorting of single cells into 96-well plates and partially automated library preparation using a liquid handling genomics robot (**Fig. 1a**). The new approach greatly enhances cell throughput, genomic coverage and contact map quality. We processed a total of 3307 single F1 hybrid 129 x Castaneus mouse ESCs with a median coverage of 182,000 distinct contacts per cell (**Fig. 1b**) using 947,000 reads per cell on average (**Extended Data Fig. 1a-c**). Analysis of various QC metrics (**Extended Data Fig. 1d-g**) confirmed the robustness of the protocol over different batches. We quantified and screened karyotypic imperfections within the single-cell sample (**Extended Data Fig. 1h**), overall retaining 2429 cells (73.4%) passing stringent quality control filters for further analysis. We found that the incidence of *trans*-chromosomal contacts in the single-cell maps (median 6.3%, **Fig. 1c**) was much lower than previously observed in ensemble Hi-C, indicative of high library quality^14^ and reflecting specific and restricted interfaces of chromosome territories (**Extended Data Fig. 2**). By pooling high quality single-cell datasets, we created an ensemble Hi-C map with reduced technical noise built upon 661 million contacts, facilitating high-resolution characterization of ensemble A/B compartments and TADs (**Extended Data Fig. 3**). More importantly, the dataset provided us with an opportunity to study the variation in chromosomal conformations across a deep sample of a dynamic nuclei pool, something that was so far impossible to assess globally and at high resolution.

**Figure 1:**
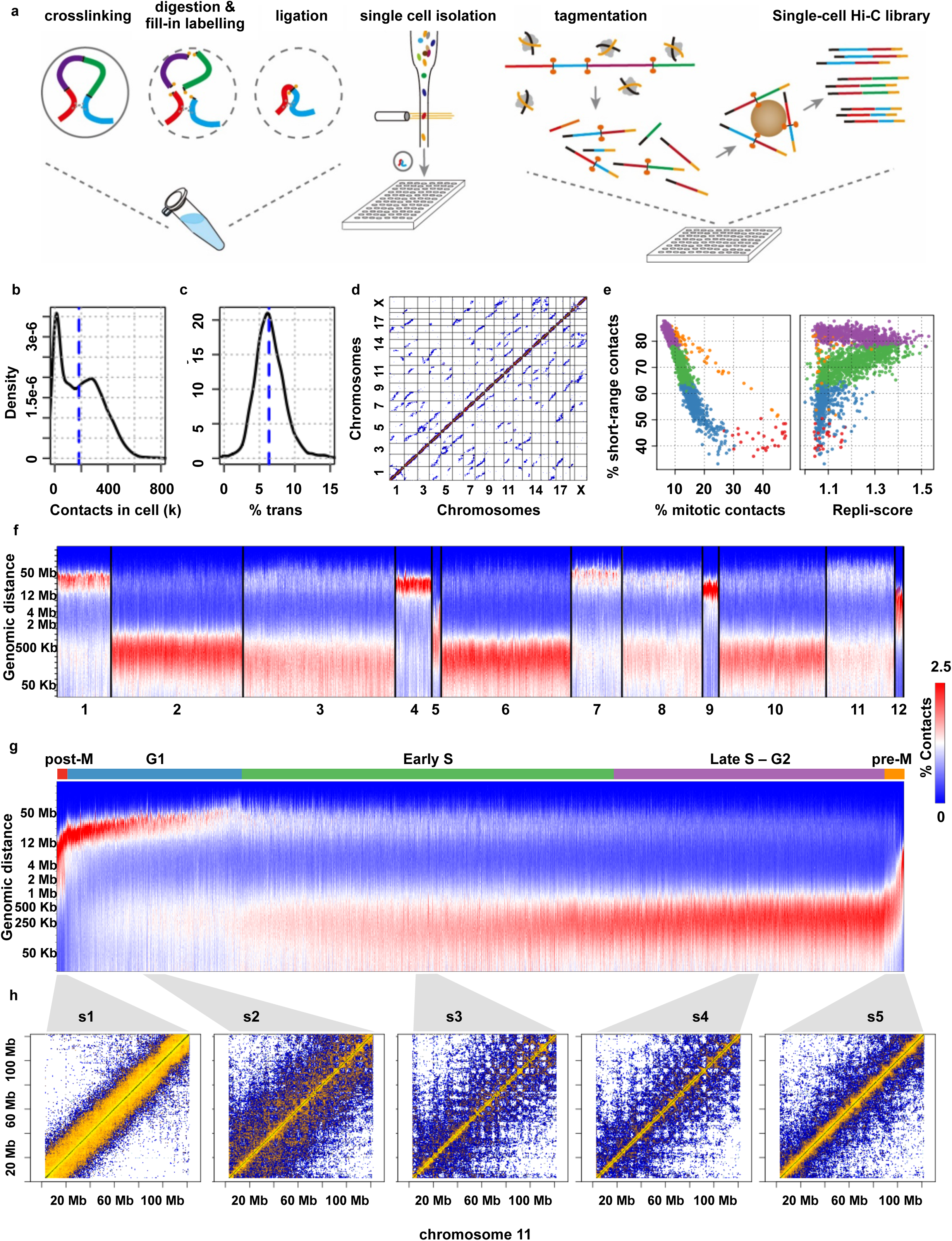
Multiplexed single-cell Hi-C reveals heterogeneous, cell cycle dependent chromosomal architectures. **a)** Single-cell Hi-C schematic. Crosslinking, restriction enzyme digestion, biotin fill-in and re-ligation are performed in nuclei with millions of cells. Individual cells are then sorted into 96-well plates for semi-automated processing to make single-cell Hi-C libraries using tagmentation, biotin pull down and PCR amplification followed by high throughput DNA sequencing. **b)** Number of contacts retrieved per cell, showing the median (∽182,000) dashed blue line. **c)** Fraction of *trans*-chromosomal contacts per cell, median (6.3%) dashed blue line. **d)** Genome-wide contact map of a representative mitotic cell (1CDX4_242) **e)** Percentage of short-range contacts (< 2 Mb) per cell compared to the percentage of contacts at the mitotic band (2-12 Mb, left) and repli-score (right). Cells are grouped by their % short-range and % mitotic contacts and coloured by group. **f)** K-means (k = 12) clusters of single-cell contact decay profiles. **g)** Single-cell decay profiles ordered by our in-silico inferred cell cycle phasing, approximate cell cycle phases are shown on top. **h)** Chromosomal contact maps (chromosome 11) of pooled consecutively ordered cells (27 cells for s1, 57 cells for s5 and 60 cells for s2- 4) marked at the bottom of **g**.

### Mitotic-like Hi-C maps

We started exploring the cell cycle dynamics revealed by single-cell Hi-C by searching for maps with mitotic signatures. Mammalian mitotic chromosomes are rod-shaped, and ensemble Hi-C on sorted M phase cells has revealed a characteristic scale of *cis*-chromosomal contact distances peaking between 2-12 Mb^12^. We examined the cells with the highest fraction of contacts at this mitotic range and found that their overall distribution of contact distances was similar to that previously observed for mitotic cells (**Extended Data Fig. 4a**). Furthermore, these single-cell mitotic maps were enriched for remarkable patterns of *trans*-chromosomal alignments (**Fig. 1d**, **Extended Data Fig. 4b**), suggesting that the chromosomes were interacting along their entire length (head-to-head and tail-to-tail) (1CDX4_242 and 1CDS2_547 in **Extended Data Fig. 4b**) as though organized by the mitotic spindle in anaphase or telophase. Other cells had similar trans-alignment patterns, but also showed evidence of head-to-tail trans-alignments (1CDX_74 and 1CDX_453 in **Extended Data Fig. 4b**), suggesting a more disordered arrangement of rod-like chromosomes, possibly indicative of cells entering mitosis or disordered chromosomal alignments at the metaphase plate. The single-cell resolution reveals details of chromosomal organisation and structure during mitosis that are not apparent in ensemble maps, and provides clear and compelling validation of the cell cycle phase of these cells. Moreover, comparison of the frequency of contacts in the mitotic range to the frequency of contacts at shorter distances (short-range contacts < 2 Mb; **Fig. 1e**, left) shows a remarkable circular pattern evocative of a gradual chromosome remodelling process as cells enter and exit mitosis. These observations suggest chromosome conformation may be used to phase single-cells at various stages of the cell cycle around mitosis.

### Single-cell replication timing

To annotate our single-cell dataset with an additional cell cycle phase, we screened for cells with evidence of DNA replication. As cells undergo genome replication in S-phase, the copy number of autosomal chromosomes increases from 2 to 4. Early-and late-replicating domains are defined by their copy number dynamics during S phase, and correlate very well with active and inactive TADs, respectively^15^. Indeed, we found that normalized TAD coverage across the single-cell dataset reflects a strong correlation between domains previously annotated as early-or late-replicating in mouse ESC^16^ (**Extended Data** **Fig. 4c-d**). We therefore computed a *repli-score* for each cell based on the copy number ratio of early-replicating regions to total coverage. G1 cells would be expected to have low repli-scores, whereas cells in early S phase would be associated with increasing scores, peaking in mid-S phase before declining again in late S phase and returning to a low level in G2/M, similar to G1. Combining repli-score with the circular pattern exhibited by the frequencies of short-range and mitotic contacts (**Fig. 1e**, left) follows exactly that expected trajectory (**Fig. 1e**, right). Repli-scores of mitotic cells are low; scores remain low as the frequency of mitotic contacts diminishes, then start to increase together with the frequency of short-range contacts, reaching a 1.5-fold enrichment peak before decreasing again to the mitotic minimal level. The repli-score suggests that the cell cycle corresponds to a clockwise transition of cells along the circular pattern exhibited by short-range and mitotic contact frequencies (**Fig. 1e**). These results link replication dynamics with a major continuous change in chromosome conformation. Together, the mitotic signatures and time of replication analysis define two major anchors to support phasing of the entire single-cell Hi-C dataset along the cell cycle.

### Integrated cell cycle phasing

As discussed above, mitotic and replicating cells show contact enrichment at specific distance scales. To generalise this observation, we clustered cells based on their contact distance distributions (**Fig. 1f**). One group of clusters (2, 3, 5, 6, 10) showed strong preference for short-range contacts, as shown for cells linked with replication above (**Extended Data Fig. 5**). A second group (clusters 1, 4, 9, 12) was characterized by sharp long-range contact peaks varying between 4 to 30 Mb (with cluster 12 including all the cells defined above as mitotic). Finally, a third group (clusters 7, 8, 11) showed intermediate distributions mixing long-range and short-range contacts, but not mitotic scale contacts. Comparative analysis of cluster parameters suggested clustering discretises a continuum of conformation changes linking the mitotic and replication states. Based on these observations, we grouped cells according to their frequencies of short-range (< 2 Mb) and mitotic (2-12 Mb) contacts (**Fig. 1e**). We first singled out cells approaching mitosis (pre-M regime, in orange) and coming out of mitosis (post-M regime, in red) by thresholding short-range and mitotic contact frequencies (see Methods). Cells in both regimes can be easily ordered by the ascending and descending frequency of mitotic contacts for the pre-M and post-M regimes, respectively. Next, we observed that cells with reduced mitotic contacts and less than 63% short-range contacts do not include the replication group and likely represent a G1 regime (blue). The mean distance of long-range contacts in these cells is correlated to the overall frequency of short-range contacts (**Extended Data Fig. 6a**), suggesting a conformation spectrum linking the mitotic state toward the replication state. Cells with over 63% short-range contacts show a continuum of repli-score and short-range contact enrichments (**Extended Data Fig. 6b**) that we used to phase single cells in two additional regimes, one based on a gradual increase in repli-score and short-range contacts (the early-S regime, in green), and one based on a gradual decrease in repli-score and further increase in short-range contacts (the late-S/G2 regime, in purple). Using these five regimes we observed a smooth transition of the distribution of genomic contact distances. The distinct patterns characterizing mitotic cells are followed by expanding long-range contacts in postulated G1 cells together with a gradual increase in short-range contacts. Short-range contacts then further increase as DNA replication progresses through the postulated early-S and late-S/G2 regimes (**Fig. 1g**, **Extended Data Fig 6c**). Since our phasing approach summarises thousands of contacts using only the contact distance distribution and the repli-score, it is highly robust to subsampling of the data. The vast majority of cells (97.75%) did not vary in their inferred cell cycle position by more than 10% when sorted in-silico using disjointed subsets of chromosomes (**Extended Data Fig. 6d**). Moreover, the contact distributions of each chromosome in each cell were consistent with the cells’ inferred cell cycle state (**Extended Data Fig. 6e**). In summary, we discovered that the global dynamics in chromosome conformation support a potential strategy for in-silico, cell cycle phasing that organises thousands of single cells along a smooth trajectory of highly-varied conformations (**Fig. 1h**) leading from the mitotic state through replication and back into mitosis.

### Cell sorting confirms inferred phasing

To confirm our Hi-C based cell cycle phasing, we analysed 1169 cells sorted based on Geminin and DNA content using fluorescence activated cell sorting (FACS) to examine how 4 distinct cell cycle populations (G1, early S, mid-S, and late-S/G2, **Fig. 2a**) are distributed in our in-silico cell cycle phasing. We found that the projected ordering of these groups was remarkably consistent with their stage as determined by FACS (**Fig. 2b**). This confirms that the Hi-C phasing strategy is compatible with coarse-grained cell cycle stages. We hypothesise that the refined ordering we impose within each stage is likely to represent cell cycle progression at high resolution. To further support this idea we analysed the distribution of key statistics for single cells ordered according to their inferred cell cycle stage. For reference, we projected the repli-score on the cell cycle progression (**Fig. 2c**), validating that it was low during G1 and G2, showed consistent gradual increase during early S phase, and decreased during late S phase. The total number of contacts (**Fig. 2d**), which unlike the repli-score is a-priori independent of the inferred phasing, was shown to increase gradually during S-phase, peaking at G2 at almost a 2-fold higher level than G1, before dropping sharply after mitosis. The fraction of *trans*-chromosomal contacts exhibits an unforeseen cell cycle dynamic, with a smooth curve from its lowest point of around 3% at G2 and mitosis, to ~8% at early S-phase (**Fig. 2e**). Next, we quantified the trans-alignment of chromosomes, which is a strong characteristic of mitotic cells. Remarkably, despite being a-priori unrelated to the inferred G1 ordering scheme, we found the trans-alignment score declined smoothly from its peak at mitosis to its nadir at the end of the G1 phase, immediately before the start of S phase (**Fig. 2f**; see **Fig. 2g** for examples). In conclusion, cell cycle phasing based on *cis*-chromosomal single-cell Hi-C contact data is corroborated by FACS to demarcate the basic stages, and is also consistent with independent continua of total number of contacts, and *trans*-chromosomal conformation metrics within each phase. Collectively, these results validate our phasing strategy and open the way to analyse cell cycle dynamics of Hi-C architectural features at high resolution.

**Figure 2:**
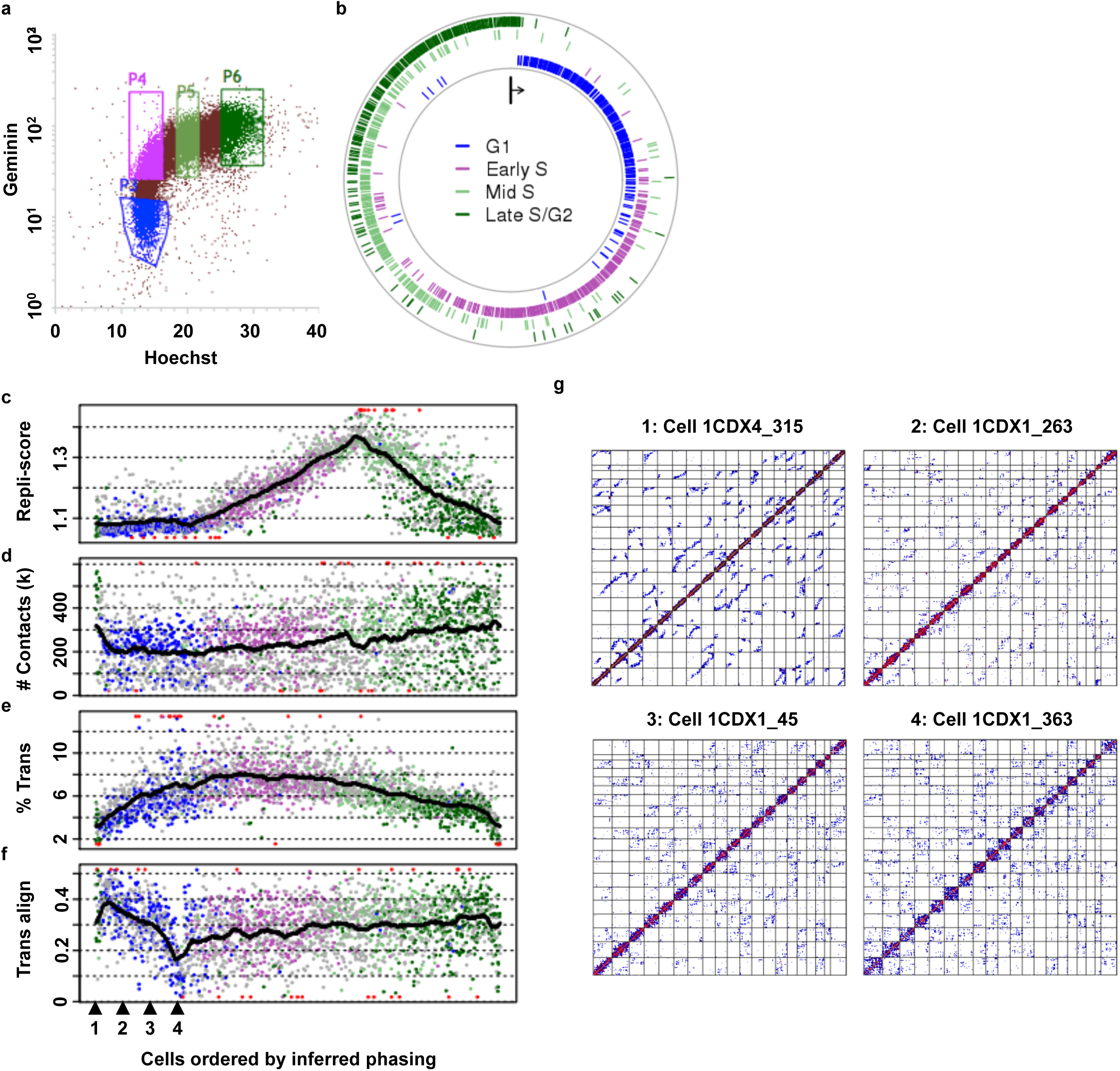
Validation of the inferred cell cycle phasing. **a)** FACS gates on Geminin and Hoechst staining used to sort cells into G1, early S, mid-S and late S/G2 phases (marked P3-P6, respectively). **b)** Positions of the FACS sorted cells along the *in-silico* phasing (cells coloured by their sorting group as in **a)**. The first position on top is the leftmost cell in **Fig. 1g**, and positions progress clockwise. **c)** Repli-score per cell ordered by the cell cycle phasing (left to right). FACS sorted cells are coloured as in **a** and **b**, the remaining cells are shown in grey. Outliers are shown in red, trimmed to the top and bottom 0.5% percentiles. Average window (n = 100) trend shown in black. **d)** Similar to **c** but showing total number of contacts per cell **e)** Similar to **c** but showing fraction of *trans*-chromosomal contacts. **f)** Similar to **c** but showing alignment score of *trans*-chromosomal contacts. **g)** Genome-wide contact maps of representative cells along the inferred G1 phase, their position marked at the bottom of **f**.

### Distinct cell cycle dynamics for TAD insulation and A/B compartment formation

We identified 2682 TAD borders in the pooled Hi-C map using analysis of contact insulation signatures^6^ (**Extended Data Fig. 3**). We created sub-pools of G1, early-S and late-S/G2 cells *in silico* to compare insulation patterns between the three major cell cycle phases (**Extended Data Fig. 7a**). We found that the locations of TAD borders are generally unchanged in these pools, but that the average intensity of the borders is dynamic (**Extended Data Fig. 7b-c**). We then calculated the mean insulation at borders for each cell (**Fig. 3a**, left) and found that the intensity of insulation varies markedly over the cell cycle. Insulation is not detectable in mitotic maps (**Extended Data Fig. 7d-e**), and emerges rapidly upon exit from mitosis in early G1. Insulation increases to its maximum during G1 but starts to decline when replication begins, until plateauing at mid S-phase at its lowest level during interphase and remaining fixed through G2 until the next cell division, when it is lost rapidly as cell approaches mitosis. We note that since insulation is defined by the depletion of short-range contacts crossing TAD borders, it may be affected by changes in the frequency of contacts in the 25 kb-2 Mb distance range. However, this overall frequency is not a good predictor for single cell insulation (**Extended Data Fig. 7f**).

**Figure 3:**
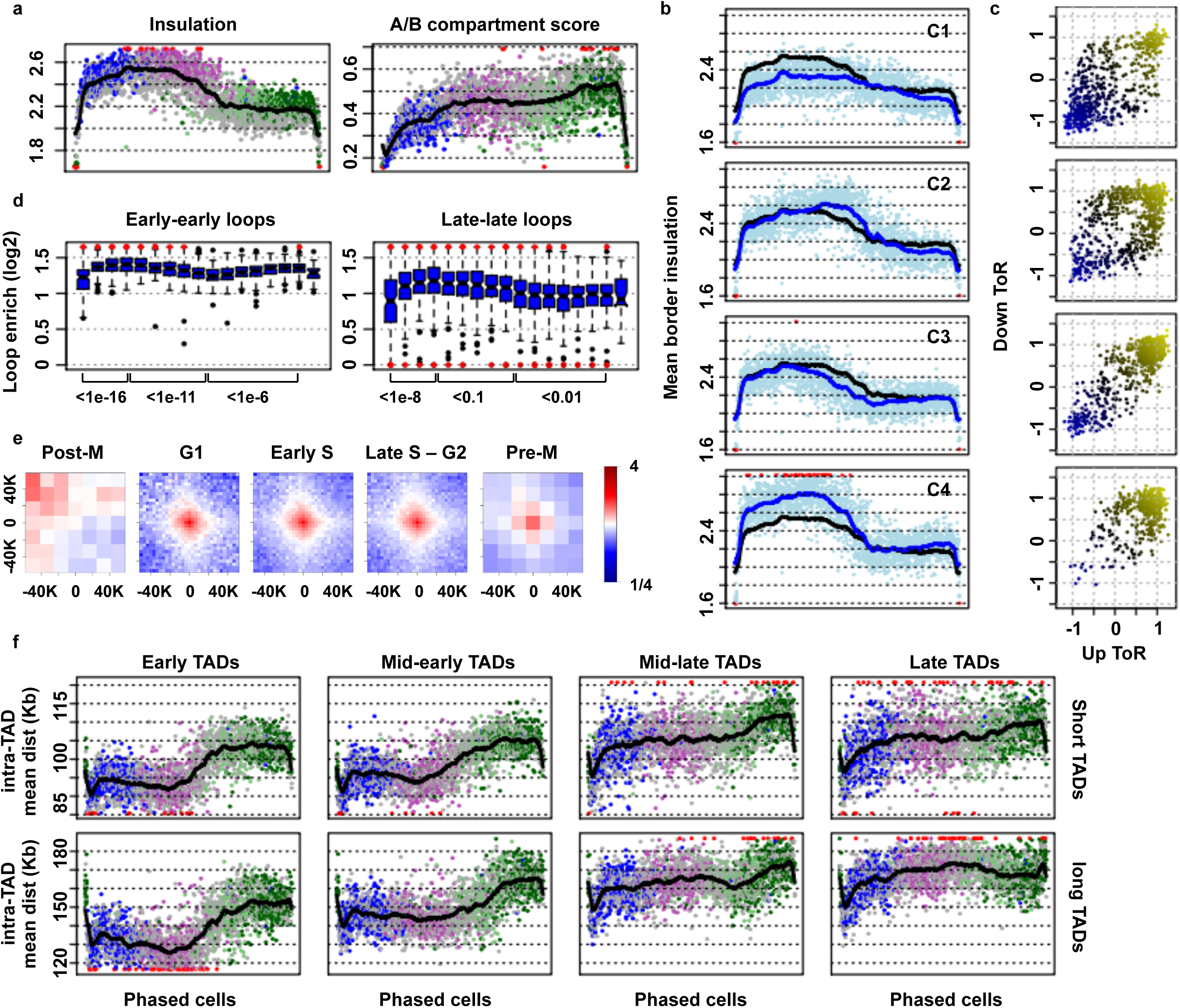
Distinct and context specific cell cycle dynamics of insulation, compartmentalization, loops, and domains condensation. **a)** Mean contact insulation on TAD borders inferred from the pooled cells (left) and A/B compartmentalization score (right) along the phased cells (display similar to **Fig. 2c**). **b**) Clustering borders (k-means, k = 4) according to their insulation profile over the inferred phasing. Showing mean insulation of the borders in a cluster (light blue dots), mean cluster insulation trend (blue line) and global trend (black line). **c)** Mean time of replication 200 kb upstream and downstream each of the **Fig. 2b** clustered borders. Early-replicating values are positive and shown in yellow, late-replicating are negative and shown in blue. **d)** Convergent CTCF loops foci enrichment (contacts concentration around a loop) over the inferred phasing, showing boxplots of 100 consecutive cells with KS-test p-values between selected groups and outliers in red. Stratifying loops by replication time of their anchors, showing early-early loops (left) and late-late loops (right) **e)** aggregated contacts around convergent CTCF loops distanced 200 kb to 1 Mb from each other, pooling cells by their *in-silico* phasing group and normalising by the expected profile using the shuffled pooled contact map. **f)** Mean intra-TAD contact distance over the inferred phasing. Stratifying domains by their length (Top: 244-490 kb, Bottom: 490-980 kb), and their mean time of replication (early, mid-early, mid-late and late). Colouring is similar to **a**.

Next, we studied the dynamics of compartmentalisation of the genome into active (A) and non-active (B) groups of TADs. We classified domains to compartments by their *trans*-chromosomal contact profile on the pooled contact map (**Extended Data Fig. 3b-d**) and then analysed the *cis*-chromosomal compartment linkage for single cells. For each cell we counted the number of A-A, B-B and A-B contacts, considering *cis*-chromosomal contacts between loci at least 2 Mb apart, and estimated the depletion of A-B contacts given the counts on A-A and B-B (Methods, **Fig. 3a**, right). Similar to insulation, compartmentalisation is weakest during mitosis, as expected from the condensed uniform structure at that stage^12^. However, in contrast to TAD insulation, compartmentalisation is weak during G1 phase and increases upon entering S phase, with a further increase in late S phase, reaching a maximum in G2, just before a rapid and complete loss when cells approach mitosis. Compartmentalisation calculated from *trans*- instead of *cis*-chromosomal contacts shows exactly the same trend (**Extended Data Fig. 7g**). The distinct cell cycle dynamics of TAD insulation and compartment structure suggests that independent mechanisms contribute to these phenomena.

### Insulation loss at TAD borders correlates with time of replication

To study cell cycle dynamics of insulation at individual TAD borders and compare it to the global trend discussed above, we grouped single cells into 24 time-slots according to their inferred cell cycle phase, and pooled data from each time point to compute an insulation time series for each border. We then clustered the borders based on their dynamics in these 24 time slots, and examined the mean insulation of borders in each cluster per cell, along the entire inferred cell cycle phasing. Borders in each cluster demonstrate similar qualitative dynamics. However, each cluster varies in timing and intensity of the deterioration in insulation during S-phase (**Fig. 3b**, shown for 4 clusters). Cluster C4 includes borders showing the strongest G1 insulation and cluster C3 borders show the earliest loss of insulation in S. Interestingly, borders in both clusters tend to be located between early-replicating TADs (**Fig. 3c**). Borders in cluster C2 show a gradual increase in insulation well into S-phase, followed by an abrupt loss of insulation in mid-S. This cluster is enriched for borders separating early-replicating TADs from mid-to late-replicating TADs (**Fig. 3c**). Cluster C1, which has the weakest G1 insulation, shows gradual and delayed loss of insulation, and is enriched for borders between late-replicating domains. These data suggest that loss of TAD insulation in S-phase is temporally coupled to the replication process. Specifically, replication through a border or in both adjacent TADs is linked to loss of insulation (cluster C1, C3-4), whereas replication of only one adjacent TAD is linked to a temporary increase in insulation (cluster C2). In all cases, insulation remains low post-replication and throughout G2, not re-emerging until the next cell cycle.

### CTCF/cohesin loops are stable over the cell cycle

We identified pairs of looping loci by statistical analysis of pooled matrices and shuffled pooled matrices (**Extended Data 7h**). We focused on 53,361 loops that were associated with convergent CTCF motifs and high probability of co-occurrence of CTCF/Cohesin (Methods). We then further grouped loops according to the replication time of their anchors (early-early vs. late-late) and computed for each group and each single cell the total number of looping contacts in 20x20 kb bins around the loop anchors, normalized by the number of contacts in the surrounding 60x60 kb bin (**Figs 3d**). We found that loop enrichment increases quickly after mitosis, is generally stable through G1, but then decreases (albeit slightly) in early S phase for early-replicating loops and in late S phase for later replicating loops. In contrast to insulation, which remains low after replication, early-replicating loops are re-intensified in late S and G2. We do not detect loops that are specific to a certain cell cycle phase (**Extended Data Fig. 7i**), suggesting that even when decreased in their average intensity during replication, loops are significantly observed in the data and do not disappear completely, as directly shown by aggregating contacts around loops from cells in each of the phasing regimes (**Fig. 3e**). However, the assay’s resolution is not sufficient to allow detection of individual loop turnover at single-cell resolution, leaving unresolved different regimes for rapid loop dynamics. Interestingly, loops are still apparent in cells approaching mitosis despite the fast chromosomal condensation, unlike cells after mitosis, which do not have any contact enrichment around loops (**Fig. 3e**). The notable lack of loop contact enrichment in mitotic maps does not necessarily imply loops are disassembled at this stage, since mitotic condensation may increase the local contact frequency around loop anchors and decrease Hi-C sensitivity at this stage.

### TAD condensation correlates with the replication process

Given the relative stability of looping contacts that demarcate TAD structures and the dynamic cell cycle rise and fall in TAD insulation, we next wished to test if chromatin structure within TADs is altered during the cell cycle, or alternatively if TADs constitute stable chromatin blocks that are re-organised dynamically during G1, S and G2. We computed average intra-TAD contact distances and used these as a measure of TAD condensation per cell, rationalizing that condensed TADs will be characterised by higher mean contact distance than de-condensed TADs. We compared TAD condensation for groups of TADs with matching sizes and variable time of replication (**Fig. 3f**). This showed that in G1, early-replicating TADs are overall de-condensed (e.g. showing lower mean intra-TAD contact distance) compared to mid-, or late replicating TADs, consistent with open chromatin in regions of high gene activity. Early-replicating TADs become even further de-condensed in early S phase, before showing a remarkable increase in condensation at mid-S, likely after their replication is complete. Following that, and during late S and G2, early-replicating TADs increase their mean condensation to levels that are almost as high as those initially observed for late replicating TADs in G1. Condensation of mid-early replicating TADs follows a similar trend with some delay and damped amplitude. Mid-late and late replicating TADs, on the other hand, show constitutively high condensation in G1 and throughout S, with further marked increases observed for mid-late TADs in G2. We note that the potential involvement of *trans*-sister chromatid contacts in driving the observed increase in mean contact distance cannot be determined unambiguously. However, the delay between replication of early domains at the start of S phase and their increased compaction in mid/late S suggests additional factors are contributing to the dynamics we observe. In summary, the data show that TADs are differentially decondensed in G1 phase consistent with their activity state and time of replication, and suggests that they are condensed in a way that temporally correlates with their replication dynamics in S phase.

### Cell cycle dynamics is recapitulated in haploid ESCs

We next generated 1257 QC positive single-cell Hi-C maps from haploid mouse ESCs, to allow systematic analysis of conformation in single copy chromosomal loci. We confirmed haploid ESC maps show comparable QC metrics to diploid cells, with less than half of the contacts per cell and some specific patterns of chromosome loss that we filtered in-silico (**Extended Data Fig. 8a-c**). We derived a pooled ensemble haploid Hi-C map and computed TAD structures, insulation and large scale A/B compartments. We then used the same approach for cell cycle phasing that was developed for diploid cells and ordered haploid cells along presumed cell cycle stages (**Fig. 4a**). This resulted in remarkably similar trends, with a typical mitotic conformation, its expansion during G1, and a marked increase in the frequency of short-range contacts during replication and G2. Repli-score for haploid cells was consistent with our ordering, as well as insulation, loop enrichment dynamics and intra-TAD condensation (**Extended Data Fig. 8d-g**). The universality of these processes shows that haploid ESCs are comparable to normal diploid cells, and a good model for studying chromosome conformation and dynamics, while providing the opportunity for unambiguous 3D modelling of single copy chromosomes for G1 cells.

**Figure 4:**
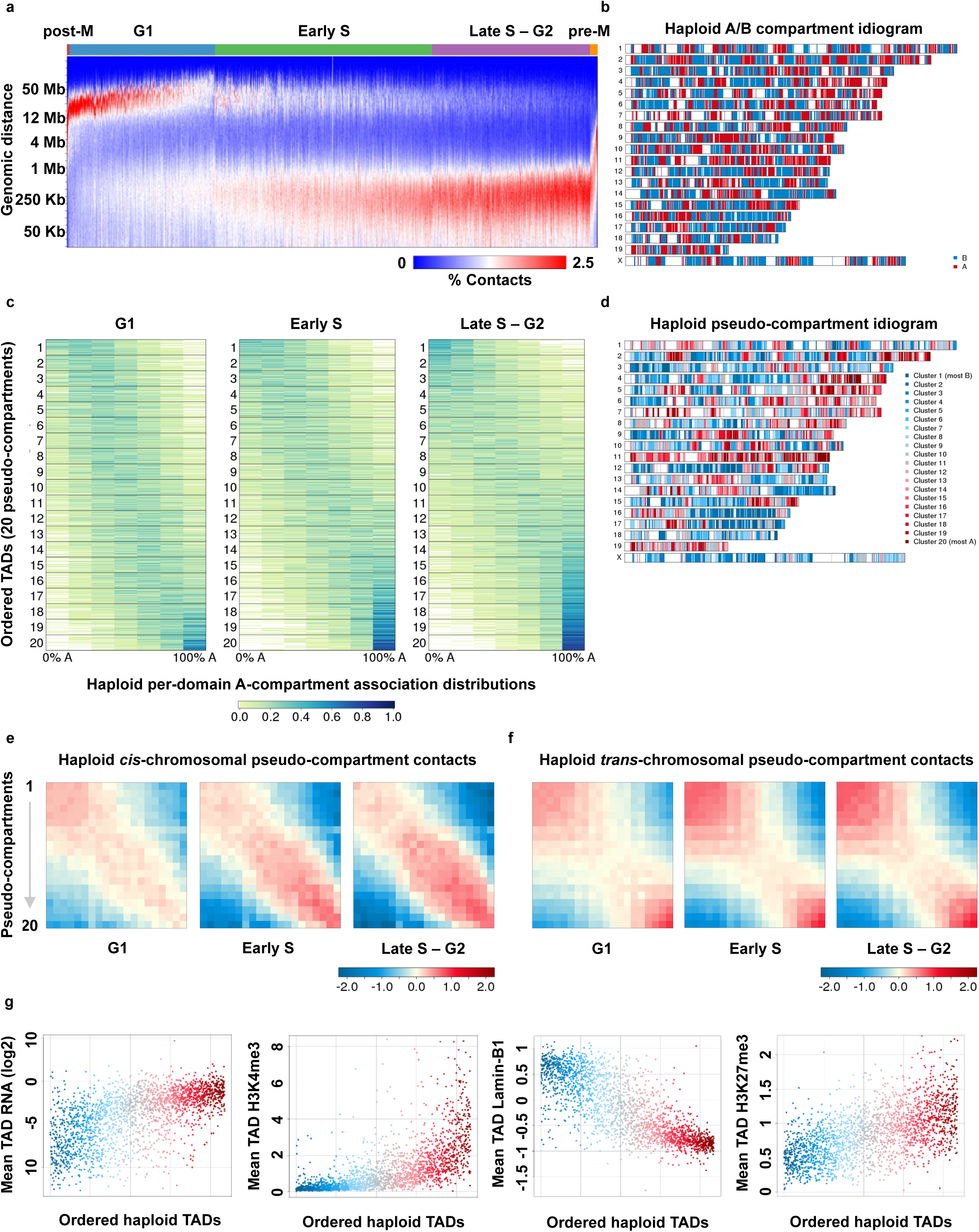
Haploid single-cell Hi-C reveals a spectrum of TAD pseudo-compartments. **a)** Cell cycle phases for single haploid ESC nuclei (x-axis ordering) were inferred using the same algorithms applied for diploid cells (**Fig. 2**). The heatmap represents the distribution of contact linear distance (y-axis, note usage of logarithmic scale). **b)** A chromosome idiogram depicts the inferred A and B compartment of each TAD, based on the pooling of haploid ESC maps. **c)** Per-domain distributions of A-compartment association across haploid cells in each of the three FAC-sorted cell populations (from left to right: G1, early S, late S to G2). For each TAD we sampled 6 long-range contacts (over 2 Mb) in each nucleus, and counted the number of A-compartment associations out of these; the distributions of a given domains percentage of A-associated contacts over the sampled cells in the three cell cycle phase groups are shown as a row in the heatmap. We used this estimated distribution of A-compartment association to sort the TADs (strongly B–associated - top, weakly A/B linked – middle, strongly A-associated – bottom, where sorting is based on the S-G2 group of nuclei). We define pseudo-compartments by splitting the set of all analysed TADs into 20 equally sized sets based on the compartment association ordering. TADs without a sufficient number of contacts (40 cells with at least 6 long-range contacts for that TAD) are not shown in the heatmap but are assigned to pseudo-compartments by majority voting and included in **d** - **g**. **d)** Colour-coded pseudo-compartments are shown on a chromosome idiogram. **e)** Long-range (> 2 Mb) *cis*-chromosomal and **f)** *trans*-chromosomal contact enrichment between TAD groups according to pseudo-compartments. **g)** TADs are sorted by their inferred A-compartment association (x-axis, left – strongly B, right – strongly A), and their mean RNA, H3K4me3, Lamin-B1 and H3K27me3 levels are shown (y-axis).

### Pseudo-compartments are defined by single-cell contact distributions

The classification of TADs into A and B compartments is established on the basis of pooled Hi-C maps (**Fig. 4b**), making it impossible to determine if individual TADs can switch compartments in specific cells or whether certain groups of cells show alternative compartment structures that converge to the average A/B model only when pooling millions of nuclei. To explore these questions at single-cell resolution we defined and computed a single-cell compartment association score for each TAD, using the number of long-range (> 2 Mb) *cis*-chromosomal contacts formed by the TAD with partner TADs in either the A or B compartments (represented in this case by early and late replicating TADs respectively). Following conservative down-sampling of the Hi-C contacts, we estimated the distributions of compartment association scores for 2175 TADs with sufficient coverage (out of a total of 2649), separately for groups of cells drawn from G1, early S and late S/G2. Clustering TADs based on these distributions (**Extended Data Fig. 9a**) showed that TADs vary quantitatively in their preferences to the A or B compartment. Despite this variation, the results also indicate that no significant group of TADs show the bimodal compartment association distribution expected to characterise domains that associate predominantly with one compartment in some cells and another compartment in other cells. The lack of bimodal TADs is particularly notable in G1 cells, for which the compartment association scores truly represent the contacts of single copy TADs. Moreover, compartment association polarizes in late S/G2, with some TADs shifting from nonspecific association in G1 or early S, to strong A- or B-association only following replication.

Interestingly, despite the simple binary nature of a two-compartment model, we observed a full spectrum of compartment association scores. This may suggest that some TADs are consistently (across all single cells) predominantly associated with either the A or the B compartment, while others are consistently associated with both, contacting A and B roughly equally in most cells. Alternatively, this data may suggest that inter-TAD contacts should be represented by a spectrum of compartments rather than a discrete classification of TADs into two or more classes. To further explore this effect we sorted TADs based on their A-compartment association in late S/G2 cells (**Fig. 4c**), deriving 20 classes of TADs that corresponded to putative *pseudo-compartments* (**Fig. 4d**). We tested how such pseudo-compartments contact one another *trans*- and *cis*-chromosomally, after normalising by expected ensemble maps to account for the genomic layout of the compartments and the increased probability of contacts in TADs that are genomically proximal. We observed significant enrichment of *cis*-chromosomal contacts between TADs within the same pseudo-compartment and within neighbouring pseudo-compartments (**Fig. 4e**). In *trans* we observed a more binary contact structure that nevertheless demonstrates enriched interaction between adjacent pseudo-compartments belonging to each of the two larger blocks (**Fig. 4f**). Together, these data support the concept that the pseudo-compartments represent a more accurate approximation of large-scale *trans*- and *cis*-chromosomal conformation than the binary notion of A (active) and B (inactive) chromosomal fractions.

### Pseudo-compartments are quantitatively characterized by epigenetic marks

To further characterize the epigenetic distributions underlying pseudo-compartments we generated RNA-seq data for haploid ES cells and studied mean RNA output per TAD as a function of pseudo-compartment ordering. We also used ChIP-seq data for histone marks from diploid mouse ESC^16^ (**Fig. 4g**), as RNA-seq data was generally consistent between haploid and diploid cells (**Extended Data Fig. 9b**). As expected, we found TADs in the strong A pseudo-compartments showed high levels of expression, while strong B TADs lacked expression almost completely. More surprisingly, we detected gradients of transcriptional activity and epigenetic marks that were quantitatively consistent with the pseudo-compartment degree of A-association (**Fig. 4g**, left). Quantitative correlation was also observed when analysing mean H3K4me3 enrichment, mean H3K27me3 levels, and mean Lamin-B association (**Fig. 4g**). Overall, the ordering of TADs according to A-association score is significantly correlated with a linear model weighting H3K4me3 and H3K27me3 scores (r=0.70, p<<0.001). In summary, the inferred pseudo-compartment structure is supported by both self-interacting contact enrichment and quantitatively variable epigenetic and transcriptional states. Whether the contact structure determines the epigenetic landscape and transcriptional control, or the epigenetic and regulatory processes drive the formation of compartment and pseudo-compartments, remains unresolved.

### Differential expansion of chromosomes and pseudo-compartments

Using single-cell Hi-C data for haploid cells that were inferred to be in G1, we developed a structure sampling approach for inferring whole genome 3D conformation. Since structure modelling is sensitive to low levels of noise, we used only high QC cells, and applied stringent contact filtering to identify contacts that are more likely to be technical artifacts (Methods), removing a median of 0.89% of the contacts from each cell (**Extended Data Fig. 10a**). We binned data at three different resolutions (1 Mb, 500 kb, 100 kb) and applied a simulated annealing protocol^11^ to sample over 30,000 whole genome 3D models for 190 single-cell datasets using the contacts as distance restraints. The resulting 3D models satisfied nearly all observed *cis*- and *trans*-chromosomal contacts (median of 0, 0.12% and 0.012% violated constraints per cell at 1 Mb, 500 kb and 100 kb resolution, respectively, **Extended Data Fig. 10b** and **c**). As few as 10,000 contacts per haploid cell produced consistent models at all resolution scales (**Extended Data Fig. 10d-e**). Cross-validation tests showed that structure modelling is sufficiently robust to re-discover *trans*-chromosomal distances between a pair of contacting chromosomes, even when all contacts between those chromosomes are discarded (**Extended Data Fig. 10f**).

To study chromosome structural changes as cells emerge from mitosis into G1, we analysed over 20,000 models from 126 cells with > 10,000 contacts per cell and < 0.5% constraint violations. Analysis of large-scale *cis*-chromosomal shapes shows that as expected, individual chromosomes rapidly decondense from a rod-like mitotic chromosome conformation to a more spherical interphase structure as G1 progresses (**Fig. 5a and b**, **Extended Data Fig. 10g** and **11a**). We monitored the expansion of chromosomes by measuring the distance between adjacent 500 kb chromatin segments in the A or B compartment in each cell (**Fig. 5c**). The A and B compartments are similarly compact upon mitotic exit (P > 0.05, two-sided KS test), but as cells progress through G1, the A compartment expands more rapidly and to a larger extent than the B compartment (P < 0.001, two-sided KS test), consistent with microscopy observations of decondensed euchromatin and compact heterochromatin. Remarkably, the twenty high-resolution pseudo-compartments we derived above generalise this observation. Decompaction of the ordered pseudo-compartments forms a gradient, with the strongest B-like pseudo-compartments being most compact and the strongest A-like pseudo-compartments being the most expanded (**Fig. 5d**).

**Figure 5:**
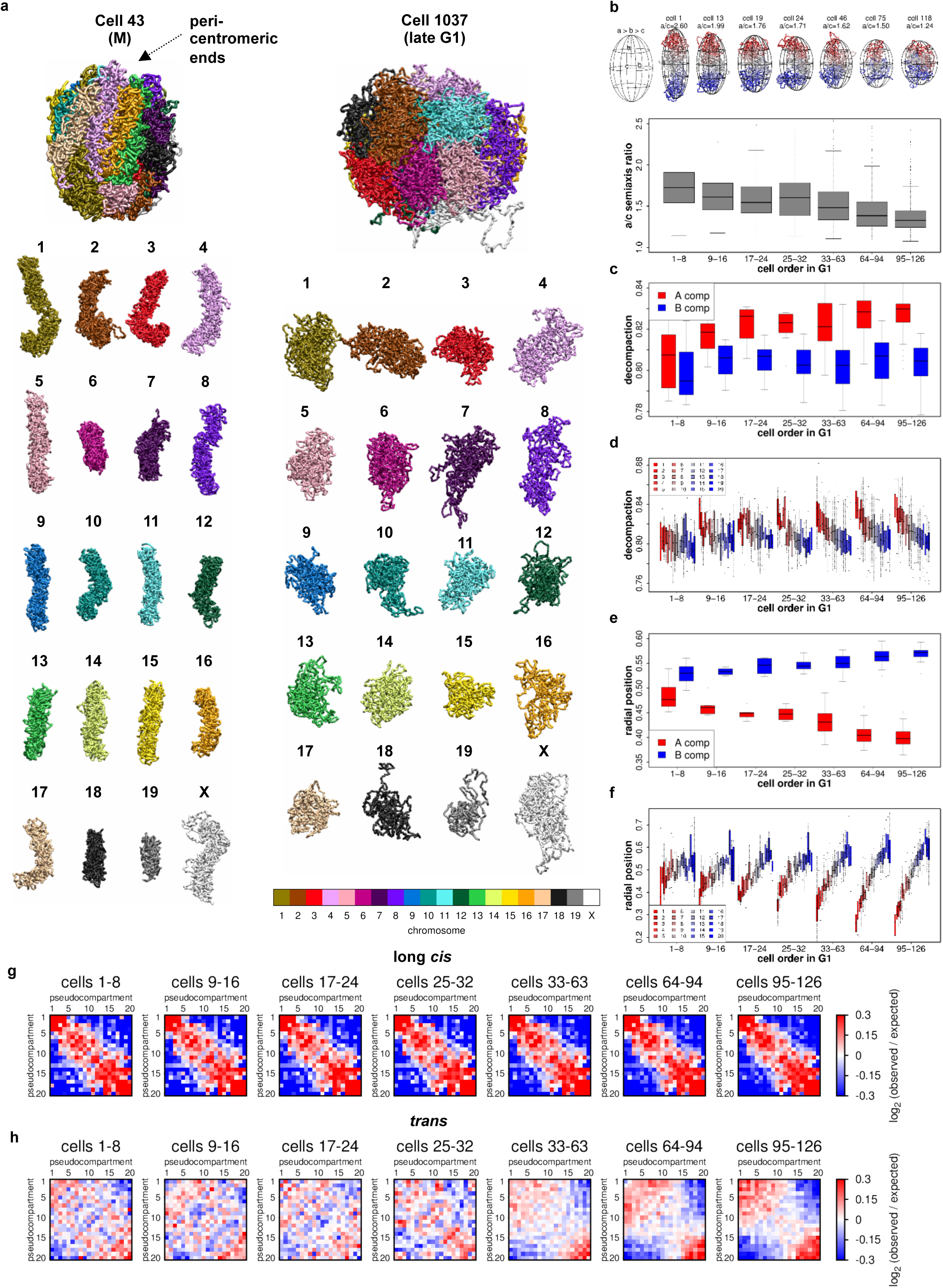
Whole genome structure modelling. **a)** Whole genome conformation for an M phase cell (NXT-43) and a late G1 cell (NXT-1037, cell 179 of the ordered 190 G1 cells), showing the whole nucleus and the separated chromosomes with peri-centromeric ends at the top. Models were computed at 100 kb resolution. **b)** Top: Chromosome 1 examples along the inferred G1 time order with their fitted ellipsoid of inertia and the inferred cell order in G1 from 1 to 126 displayed. The longest-to-shortest (a/c) semiaxis ratio is an indicator of how elongated the chromosome's shape is. Bottom: Chromosome decondensation in G1 is illustrated by the distribution of the a/c semiaxis ratio of all chromosomes of cells in 7 time groups along the inferred G1 time order. **c)** Decompaction of chromatin in the A and B compartments along the inferred G1 time order, defined as the average distance between neighbouring 500 kb segments of the same compartment. **d)** Same as c for the 20 pseudo-compartments. **e)** The average radial position of the A and B compartments within the nucleus along the inferred G1 time order, defined as the volume fraction enclosed by a sphere of that radius. Zero corresponds to the nucleus centre, 1 corresponds to the peripheral end. **f)** Same as e for the 20 pseudo-compartments. **g)** Long-range (> 2 Mb) cis-contact enrichment between the 20 pseudo-compartments. As control, contacts were shuffled uniformly keeping the same number of contacts for each compartment (see Methods). **h)** Same as **g** for *trans*-contact enrichment between the 20 pseudo-compartments.

We then computed the mean radial positioning of the A and B compartments as cells exit mitosis. Radial positions are similar in early G1 cell models, but differences soon emerge and strengthen as G1 progresses, with B compartments occupying more peripheral locations and A compartments tending toward the nuclear interior (P < 0.001 from group 2, two-sided KS test, **Fig. 5e**). The radial positioning of pseudo-compartments exhibits a striking trend that expands this concept, revealing a clear gradient of the most B-like pseudo-compartments in the periphery to the most A-like pseudo-compartments occupying more internal locations as G1 progresses (**Fig. 5f**). The structural models also confirm and refine the preferential contacts among pseudo-compartments (as shown in **Fig. 4e**,**f**), and show that long-range *cis*-contacts are reestablished very early in G1 phase while trans compartmentalisation is not fully reestablished until later in G1 (**Fig. 5g**,**h**). These findings are consistent with previous 4C observations identifying long-range *cis*-contacts occurring in early G1 between TADs with similar compartment status and linking those contacts to the establishment of the replication timing decision point^13^.

Finally, we looked at spatial clustering between peri-centromeric regions of chromosomes with Nucleolar Organiser Regions^17^ (NOR) at their acrocentric tips (chromosomes 11, 12, 15, 16, 18 and 19). The NORs on these chromosomes contribute to the formation of nucleoli, which could result in non-random spatial clustering of their centromeric and peri-centromeric regions. In early G1, pericentromeric regions of NOR or non-NOR chromosomes are similarly close to one another (P > 0.5 two-sided KS test), but as G1 progresses, non-NOR peri-centromeric regions become more distant from one another while NOR chromosome pericentromeric regions remain close (P < 0.01 two-sided KS test, **Extended Data Fig. 11b**), consistent with organization of NOR chromosomes around nucleoli. We conclude that our structural modelling approach validates key observations of our statistical analyses of single-cell Hi-C data and provides additional metrics to track genome organisation from late mitosis through G1 phase.

## DISCUSSION

Using new high throughput single-cell Hi-C technology, we have generated and analysed single-cell contact maps capturing the chromosomal conformations of thousands of diploid and haploid mouse ESCs. We confirmed using statistical analyses and whole genome structural modelling of single-copy haploid G1 cells that genome folding is highly heterogeneous at the single-cell level, such that average contact mapping using standard Hi-C may blur together remarkably different structures. The variation in genome folding, however, is far from random, and reflects a combination of deterministic dynamics and stochastic effects. Our data suggest that the most dramatic source of deterministic dynamics in the mouse ESC system is the cell cycle. The cell cycle dynamics of chromosome folding is known to accommodate condensation for mitosis and subsequent de-condensation in G1, but it has been so far unclear how chromosomes re-establish and stabilise their interphase conformation following mitosis. In addition, the global conformational changes associated with the replication process have so far only been characterised for specific loci using bulk analysis of synchronized cells^13^. Our data show that chromosome folding in mouse ESCs undergoes at least three major structural transitions during the cell cycle; condensation in preparation for mitosis, a rapid expansion in early G1, and extensive remodelling during replication. The effects of these major transitions extend throughout the entire cell cycle, such that the genome is not stably folded at any point in the cycle. The chromosome dynamics of cells with slower cycles, or post-mitotic cells, remain to be explored.

TADs have previously been identified as a key feature in ensemble Hi-C maps. TADs provide potential intermediate organizational units that package several genes and regulatory elements, but not entire chromosomes^3–5^. The presence of a TAD in a Hi-C matrix can represent a mixture of distinct mechanisms that together can contribute to the observed insulation between adjacent TADs. One type of TAD-promoting mechanism promotes contacts within TADs or at their borders, such as by long-range looping contacts. Other mechanisms that decrease the probability of contacts across TAD borders can be independent of specific loops, and involve differential domain activity (transcription), differential domain time of replication or differential domain tethering (to the nuclear lamina or other elements). Cell cycle phasing of single-cell Hi-C maps shows that overall, insulation peaks at G1 and decreases substantially during replication. The reduction in total contact insulation at individual borders is temporally coupled to the time of replication of the genomic domains around it. Importantly, when total insulation is high in G1, TADs form substantially more long-range contacts with non-adjacent TADs within their chromosomal territories. This may imply that despite showing strong TAD structure in G1, domains may be similarly or even less insulated from the rest of the chromosome in G1 compared to S/G2. The reduction in TAD insulation during and following replication may reflect a general reduction in the flexibility of chromosomal territories, possibly in part due to packaging of sister chromatids, which are being tightened up and re-organised in anticipation of later mitosis.

In contrast to contact insulation, the enrichment of contacts at the loops that pair borders of most mammalian TADs is generally stable throughout the cell cycle. A possible exception to this is mitosis, but it is unclear if Hi-C analysis, even at the single-cell level, is sensitive enough to determine conservation of mitotic loops or lack thereof, given a likely increase in competitive ligation of chromosomal elements in the compact mitotic chromosomes. The stability of loops does not imply rigidity of internal TAD structure. The data suggest active, early-replicating TADs are decondensed in G1 and early S compared to inactive, late-replicating TADs, but become significantly more condensed after replication. This further supports the idea that chromosomal conformation post replication is organised more tightly and compactly, well before mitosis takes place.

The demarcation of the genome into two nuclear compartments (known as A and B) was previously observed in ensemble maps in a way that strongly correlates with transcriptional activity^2^. It was also shown that TADs in the B compartment are late replicating and anchored to the nuclear lamina^15^, and that further epigenetic analysis may dissect chromosomes into more than two compartments^7,18,19^. Single-cell Hi-C in haploid ESCs allows for the first time the complete structural modelling of whole genomes. The resulting 3D models confirm statistical observations that show the association of TADs with nuclear compartments can be refined significantly. The data support organisation of TADs along a spectrum of preferred radial positions going from interior and highly active, through intermediate radial position and intermediate activity, to highly peripheral and inactive. The discovery of a pseudo-compartment spectrum generalises but does not contradict the previous two-compartment model, and shows that TADs are committed to their pseudo-compartment in a nearly deterministic fashion. We could not find TADs that associate with variable pseudo-compartments in different cells, although the nuclear localisation of each domain was shown to be dynamic during the cell cycle. This surprising observation shows that despite the high variability of chromosomal conformation at the single-cell level, some key architectural properties of the genome are maintained with minimal tolerance.

Importantly, single-cell Hi-C can now allow high resolution analysis of other factors driving chromosome folding and genome regulation, as we demonstrated here for Nucleolar Organizing Regions. While many other aspects of dynamic nuclear organization can now be explored at high resolution using single-cell approaches, it will become extremely important to interpret both single-cell studies and ensemble Hi-C experiments through the cell cycle prism introduced here. Focusing experimentally on G1-sorted populations represents one partial solution for controlling cell cycle confounders on Hi-C studies, but as shown here, G1 nuclei populations are still extremely dynamic. An emerging challenge is therefore the derivation of comprehensive models, combining cell cycle chromosomal dynamics with gene regulatory processes during differentiation or response to stimuli. Using such models, it may become possible to understand how nuclei are capable of supporting the remarkably deterministic and pronounced chromosomal dynamics of mitosis and replication, while implementing robust and flexible gene regulatory programs, a question that is rapidly becoming one of the key challenges in epigenetics.

## Methods

Manuscript references^1–19^

### Cell culture, fixation and sorting

Two mouse ESC lines were used. One diploid F1 hybrid ESC from an intercross of Mus musculus 129/SV-Jae and Mus musculus castaneus (129 × Castenious,a gift from Joost Gribnau) and one haploid ESC^20^ (H129-1; ECACC 14040203) were grown either in 2i medium without feeder cells or ES-DMEM with fetal bovine serum (FBS) on feeder cells^21^. For harvesting ESCs from cultures on feeder cells, the feeder cells were depleted using Feeder Removal MicroBeads (Miltenyi Biotec), followed by fixation and permeabilisation of the enriched ESCs and staining for Oct-3/4- immunoreactivity as described below, and Oct-3/4-positive cells were collected by Fluorescence-activated cell sorting (FACS), were ready for subsequent Hi-C processing.

To fix the cells, 50 to 200 million ESCs were suspended in relevant medium and fixed for 10 min by adding formaldehyde at a final concentration of 2% at room temperature before quenching with 127 mM glycine for 5 min on ice. The cells were washed with PBS and permeabilised in 10 mM Tris-HCl (pH 8), 10 mM NaCl, 0.2% IGEPAL CA-630 with cOmplete EDTA-free protease inhibitor cocktail (Roche) for 30 min on ice with intermittent agitation, and spun to collect the nuclei pellet, which was ready for Hi-C processing.

To enrich for specific cell types before the Hi-C process, cells were blocked with PBS-FT (5% FBS, 0.1% Tween-20 in PBS) at room temperature for 1 hr, incubated with a primary antibody in PBS-FT as required (anti-Oct3/4, Santa-Cruz sc-5279, at 1:100 dilution; or anti-Geminin, Abcam 175799 at 1:400 dilution) for 1 hr on ice. Samples were then washed with PBS-FT, and incubated with the secondary antibodies (Alexa Fluor 647-anti-mouse IgG, Thermo Fisher A-31571, or Alexa Fluor 555-anti-rabbit IgG, Thermo Fisher A-21428) for 30 min on ice, washed with 2% FBS in PBS, stained with Hoechst 33342 at the final concentration of 15 ?g/ml and subjected to FACS by Influx (BD Biosciences) to collect nuclei cells of interest.

### Hi-C processing

The cells (approximately half million to several million) were washed with 1.24 × NEBuffer 3 (New England Biolabs; 62 mM Tris-HCl [pH 7.9], 124 mM NaCl, 12.4 mM MgCl_2_, 1.24 mM DTT) and suspended in 400 μl of 1.24 × NEBuffer 3. Six μl of 20% SDS was added and incubated at 37°C for 60 min with constant agitation, then 40 μl of 20% Triton X-100 was added and incubated at 37°C for 60 min with constant agitation. Next, 50 μl of 25 U/μl Mbo I (New England Biolabs) was added and incubated at 37°C overnight with constant agitation. To label the digested DNA ends, 1.56 μl of 10 mM dCTP, 1.56 μl of 10 mM dGTP, 1.56 μl of 10 mM dTTP, 39 μl of 0.4 mM biotin-14-dATP (Thermo Fisher) and 10.4 μl of 5 U/μl DNA polymerase I, large (Klenow) fragment (New England Biolabs) were added and incubated at 37°C for 45 min with occasional mixing. The sample was then spun and supernatant partially removed leaving 50 μl with cells, followed by addition of 100 μl of 10x T4 DNA ligase reaction buffer (New England Biolabs), 10 μl of 100x BSA (New England Biolabs), water and 10 μl of 1 U/μl T4 DNA ligase (Thermo Fisher) were added to make the total volume 1 ml, and incubated at 16°C overnight. Then the nuclei were passed through a 30 μm cell strainer and single nuclei were sorted into individual empty wells in 96 well μl ates using an Influx cell sorter. The μl ates were sealed and stored at - 80°C until further processing.

### Single-cell Hi-C library preparation

To prepare single-cell Hi-C libraries from single nuclei in 96 well μl ate, 5 μl of PBS was added to each well, the μl ate was sealed and crosslinks reversed by incubating at 65°C overnight. Hi-C concatemer DNA was fragmented and linked with sequencing adapters using the Nextera XT DNA Library Preparation Kit (Illumina), by adding 10 μl of Tagment DNA Buffer and 5 μl of Amplicon Tagment Mix, incubating at 55°C for 5 min, then cooling down to 10°C, followed by addition 5 μl of Neutralize Tagment Buffer and incubation for 5 min at room temperature. Hi-C ligation junctions were then captured by Dynabeads M-280 streptavidin beads (Thermo Fisher; 20 μl of original suspension per single-cell sample). Beads were prepared by washing with 1 × BW buffer (5 mM Tris-Cl pH 7.5, 0.5 mM EDTA, 1 M NaCl), resuspended in 4 × BW buffer (20 mM Tris-Cl pH 7.5, 2 mM EDTA, 4 M NaCl; 8 μl per sample), and then mixed with the 25 μl sample and incubated at room temperature overnight with gentle agitation. Using a Bravo automated liquid handling system (Agilent Technologies), the beads were then washed four times with 200 μl of 1 × BW buffer, twice with 200 μl of 10 mM Tris-Cl pH 7.5 at room temperature, and resuspended in 25 μl of 10 mM Tris-Cl pH 7.5. Single-cell Hi-C libraries were amplified from the beads by adding 15 μl of Nextera PCR Master Mix, 5 μl of Index 1 primer of choice and 5 μl of Index 2 primer of choice. Samples were then incubated at 72°C for 3 min, 95°C for 30 sec followed by the thermal cycling at 95°C for 10 sec, 55°C for 30 sec and 72°C for 30 sec for 12 or 18 cycles, then incubated at 72°C for 5 min. The supernatant was separated from the beads, and the 96 supernatants from a 96 well μl ate that had 12 cycles of amplification were combined together whereas the supernatants from 18 cycles of amplification were processed uncombined. The combined or uncombined supernatant was purified with AMPure XP beads (Beckman Coulter; 0.6 times volume of the supernatant) according to manufacturer's instructions and eluted with 10 mM Tris-Cl pH 8.5 (100 μl when 96 samples were combined; 30 μl when sample was uncombined). The eluate was purified once more with AMPure XP beads (equal volume to the previous eluate) and eluted with 11 μl of 10 mM Tris-Cl pH 8.5.

### Strand-specific nuclear RNA-seq library preparation

Ten to 20 million unfixed ESCs (grown in 2i medium without feeder cells) were washed with cold PBS, resuspended in 0.5 ml of cold buffer RLN (50 mM Tris pH 7.5, 140 mM NaCl, 1.5 mM MgCl_2_, 1 mM DTT, 0.4% IGEPAL CA-630) and incubated for 5 min on ice. The sample es were spun and the resultant pellets (nuclei) were washed with buffer RLN and subject to RNA isolation using RNeasy Mini Kit (Qiagen) and QIAshredder spin columns (Qiagen) according to the instructions from the manufacturer. The strand-specific RNA-seq libraries were prepared from 200 ng each of nuclear RNA samples using TruSeq Stranded Total RNA LT Sample Prep Kit (Illumina) according to manufacturer's instructions.

### library QC and sequencing

Before sequencing, the libraries were quantified by qPCR (Kapa Biosystems) and the size distribution was assessed with Agilent 2100 Bioanalyzer (Agilent Technologies). They were sequenced by either 2 × 50 bp, 2 × 75 bp or 2 × 150 bp paired-end run by HiSeq 1500, HiSeq 2500 or NextSeq 500 (Illumina).

### Sequence processing pipeline

Paired-end reads of sequencing batches were de-multiplexed to single cell datasets based on the two 8 bp unique identification tags attached to each cell during the Tn5- transposase tagmentation. Reads lacking a perfect match were discarded. Reads were broken down to segments on their matches to the MboI recognition site (GATC), concatenating segments shorter than 16 bp with their adjacent segment. Each segment was independently mapped to the *Mus Musculus* genome (assembly mm9) Using Bowtie2^22^ in *end-to-end* alignment mode. The uniquely mapped (MAPQ > 36) segments from each paired-end read were consolidated into a chain of segments by merging segments mapped to overlapping genomic regions. Next, the segments chain was translated to a chain of fragment-ends (fends) by associating each segment with its downstream fend. Finally, adjacent fend pairs from all chains were used to comprise the contact map of the cell. All the analysis following used the distinct contacts, ignoring the number of times each contact appeared in the map.

### Quality controls and selection of high quality contact maps

We calculated several metrics to evaluate the quality of each contact map:

1. *Coverage:* Total number of contacts
2. *Trans fraction:* Fraction of inter-chromosomal contacts out of the total number of contacts
3. *Non-digested fraction:* fraction of contacts between fends distanced less than 1 kb from each other out of the total number of contacts. The vast majority of these ultra-close cis contacts originate from non-digested DNA and therefore this metric quantifies the digestion efficiency of the experiment.
4. *Max chromosomal coverage aberration:* We calculated coverage enrichment per chromosome and cell by comparing the number of contacts a chromosome forms with the expected number given the mean fraction of contacts per chromosome across the entire single cell pool. The *Max chromosomal coverage aberration* of a cell is the maximal enrichment (or depletion) of this log-ratio across all its chromosomes.

The following filters were applied on the contact maps to select high quality ones:

| Metric | Diploid condition | Haploid condition |
| --- | --- | --- |
| Coverage | 20K-700K | 10K-1M |
| Trans fraction | < 15% | < 20% |
| Non-digested fraction | < 55% | < 50% |
| Max chromosomal coverage aberration | ½ - 2 | ½ - 2 |

We found no systematic technical difference between 2i- and serum- cultured cells (**Extended Data Fig. 1f-g**), so we applied the same quality control filters for both and analyzed them together.

In our previous version of the single-cell Hi-C protocol^11^ we found that contacts supported by a single read (singleton contacts) represented by large spurious ligations and were therefore discarded. In contrast, the improved single cell Hi-C approach shows no significant difference in QC metrics between singly and doubly covered contacts. For example, we found that singleton contacts have the same cis to trans ratio as multiply covered contacts (mean of 93% versus 92.9%), suggesting low level of spurious contacts is represented in both classes. We note that in addition to overall improved processing robustness and quality, the sequencing depth of our current libraries, in contrast to the previous version, is far from saturation (**Extended Data Fig. 1b-c**) so the fraction of singleton contacts is much higher (mean 25%, **Extended Data Fig. 1a**).

### Repli-score and time of replication analysis

Repli-score quantifies the coverage enrichment of early replicating regions over late replicating ones. To calculate it we characterized each fend by the mean Repli-chip value around it (10Kb upstream and downstream), using replicate 1 of the *Mus Musculus* 129 ES-D3 Repli-chip dataset^16^. Repli-chip values are bimodal around zero, with early replicating loci having positive values and late loci negative ones. The repli-score of a cell is the frequency of early replicating fends in its contact map, divided by the minimal frequency found across all cells (to scale the score to be relative to one).

The generation of ordered correlation matrix of normalized domain copy-number (**Extended Data Fig. 4c**) is very sensitive to chromosomal copy-number variations. Therefore we selected cells with strictly normal chromosomal coverage (cells with *Max chromosomal coverage aberration* between 0.82–1.23), ending up using 51% of the cells. We next counted per cell the total number of contacts formed by each domain (*trans* and *cis* above 23.17Kb). Domains without time of replication information and domains with extremely low (< 0.04) or high (> 0.16) mean number of contacts per Kb (total of 4.2% of the domains) were discarded. The 50 domains with the earliest mean time of replication and the 50 domains with the latest one were marked as top-early and top-late, respectively. Contact numbers were transformed to contact frequencies. The pearson correlation matrix between all pairs of domains was then calculated, and domains were ordered by their coverage profile mean pearson correlation with the top-early domains coverage profile minus the mean pearson correlation with the top-late profiles.

### In-silico cell phasing over the cell cycle

We counted the number of *cis* (intra-chromosomal) contacts per cell, binning contacts by distance into logarithmic bins (143 bins, first one for contacts distanced < 1Kb, then each bin covers an exponent step of 0.125, using base 2). Contacts in bins 1–37 were found to be noisy and were discarded, making bins 38–143 the valid bins. The following metrics were used to phase the cells:

- %*near* – percentage of contacts in bins 38-89 out of all valid bins
- *%mitotic* – percentage of contacts in bins 90-109 out of all valid bins
- *farAvgDist* – mean contact distance considering bins >= 98
- *rawRepliScore* – fraction of early-replicating fends out of all fends in the contact map (see above for details)

Each cell was assigned to a group and cells within each group were ordered by these criteria (scale means subtracting the mean and dividing by the standard deviation):

| Group | Typical phase | Assignment Criteria |  | Ordering by |
| --- | --- | --- | --- | --- |
|  |  | Diploid cells | Haploid cells |  |
| 1 | Post-M | $\% \text{ mitotic} \geq 30 \wedge \% \text{ near} \leq 50$ | $\% \text{ mitotic} \geq 30 \wedge \% \text{ near} \leq 42$ | $-\% \text{ mitotic}$ |
| 2 | G1 | $\% \text{ near} \leq 63$ | $\% \text{ near} \leq 61.1$ | $\text{scale}(\% \text{ near}) + \text{scale}(\text{farAvgDist})$ |
| 3 | Early to mid-S | $63 < \% \text{ near} \leq 78.5$ | $61.1 < \% \text{ near} \leq 77$ | $\frac{\% \text{ near}}{\text{VAR}(\% \text{ near})} + \frac{\text{rawRepliScore}}{\text{VAR}(\text{rawRepliScore})}$ |
| 4 | Mid-S to G2 | $\% \text{ near} > 78.5$ | $\% \text{ near} > 77$ | $\frac{\% \text{ near}}{\text{VAR}(\% \text{ near})} - \frac{\text{rawRepliScore}}{\text{VAR}(\text{rawRepliScore})}$ |
| 5 | Pre-M | $\% \text{ near} > 50 \wedge \% \text{ near} + 1.8 \times \% \text{ mitotic} > 100$ | $\% \text{ near} > 42 \wedge \% \text{ near} + 1.8 \times \% \text{ mitotic} > 100$ | $\% \text{ mitotic}$ |

Cells are first assigned to groups 1 and 5 and those remaining are assigned to groups 2-4. To compensate for the arbitrary thresholds (which may introduce discontinuity on the interfaces between the three main phases), the initial assignment to groups is then fine-tuned by clustering the contact distance distribution profiles of the cells to small clusters (k-means, k=number of cells/10) and reassigning cells in each cluster to the majority vote of that cluster. This resulted in reassigning 5% and 10% of the cells in the diploid and haploid datasets, respectively, with all reassignments involving adjacent groups.

To test the stability of our phasing procedure among the different chromosomes we compared the positions of cells by phasing using disjoint sets of chromosomes with roughly the equivalent length (**Extended Data Fig. 6d**, 1^st^ set: chromosomes 2, 3, 5, 6, 9, 10, 12, 15, 17 and 19, 2^nd^ set: chromosomes X, 1, 4, 7, 8, 11, 13, 14, 16, and 18). Phasing positions determined by both sets largely concur, with most of the differences concentrated on the interfaces between G1 and early S, or early S late S/G2 groups. The positions of only 54 out of the 2393 cells (2.25%) differ by more than 10%.

### Pooled contact maps

Contacts from single cell maps were pooled together to generate aggregated maps. We used the total number of contacts when creating ensemble-like maps, e.g. for the pool of all cells and the phase groups, and distinct contacts when creating contact maps comprised from a few dozens of cells (e.g. **Fig 1h**, **Fig 3b**). To normalize the pooled maps (**Extended Data Figs. 3a** and **7h**), and to compare pooled contacts around epigenetic landmarks (**Fig. 3e**) we sampled a randomized ensemble Hi-C map^23^ and compared contact distributions in the observed and randomized data using the Shaman tool (https://bitbucket.org/tanaylab/shaman). Briefly, we shuffled contacts using a Markov Chain Monte Carlo-like approach, first within each chromosome and then genome-wide, such that the primary factors that define contact distributions are preserved - the marginal coverage and contact distance distribution - but any compartment or TAD structure that may be present is not retained. The resulting expected ensemble map has an identical number of contacts for each locus, and identical probability for a contact at a given genomic distance, as in the observed map. We generated global shuffled maps, as well as maps for each cell-cycle phase.

### Insulation, domain and border calling and domains initial A/B classification

The insulation of a locus (**Supplementary Fig. 1**) corresponds to depletion of contacts in the ‘a’ region. We calculate an insulation score by considering a distance around the locus (scale) and comparing the total number of contacts that violate insulation (in region a) with the total number of contacts around the locus (in regions a, *b_1_* and *b_2_*). In both counts contacts distanced less than 1Kb from the diagonal are ignored (these are mostly non-digested contacts). The negative log-ratio of these counts correlates positively with strong insulation. We calculated insulation on the pooled contact map in 1Kb resolution, using a scale of 300Kb. Domains were then identified as contiguous regions in which insulation score is below the 90% quantile of the genome-wide distribution. Domains shorter than 20Kb or longer than 4Mb and domains containing a non-mappable region longer than 25kb were discarded, resulting in 2691 domains (or 2709 domains for the haploid pool of cells).

**Supplementary Figure 1:**
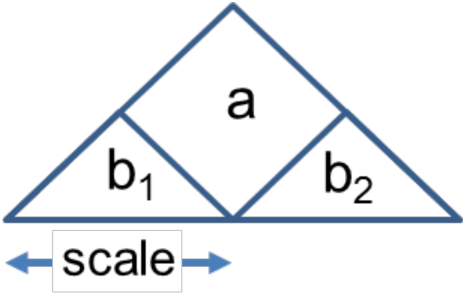
Locus insulation.

The initial assignment of domains to the A- and B- compartments was done by k-means clustering (K=2) the inter-domain contact profile, using only interchromosomal *(trans)* contacts, and computing log2 ratio of observed and expected (based on genomic length) inter-TAD contacts^18^. Domains in each cluster exhibit distinct epigenetic and time of replication signatures (**Extended Data Fig. 3c-d**) pointing us to assign the 1058 domains in cluster 1 to the B-, inactive-compartment and 1632 domains in cluster 2 to the A-, active compartment

Domain borders were called from the insulation profile of the pooled map by identifying highly insulating regions between domains (insulation above the 90% quantile) and selecting in each element the 1KB with highest insulation score. To calculate the cell mean insulation over a set of borders *B* (either all borders as in **Fig. 3a** or a subset of borders as in **Fig. 3b**) the total number of border violating contacts is compared to the total number of contacts around the border. Denoting *A*[*b*] as the number of contacts in the *a* region of border *b* (**Supplementary Fig. 1**) and similarly *B_1_*[*b*] and *B_2_*[*b*] (see schematic figure above):

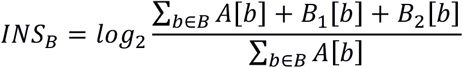

To cluster borders by their insulation profile across the inferred cells phasing (**Fig. 3b**) cells were divided to 24 slices (100 cells in each slice grouped according to phasing), insulation was calculated per border and slice and the resulting 24 length border insulation profiles were clustered (k=4). Fig 3b shows per cell (and not per slice) the mean insulation over each of the resulting clusters.

### Inter-chromosomal alignment

The head-to-head alignment of chromosomes in single-cell mitotic Hi-C maps, likely represents anaphase or early telophase nuclei (**Extended Data Fig. 4b**). To quantify the degree of head-to-head alignment of the chromosomes of each cell, we extracted trans-chromosomal contacts and scaled the coordinates of the contacting fends by the length of their respective chromosomes. Alignment was then approximated using the Pearson correlation between the two scaled fend coordinate vectors (**Fig. 2f**).

### A/B compartment score

Intra-chromosomal, Inter-domain contacts were classified according to the compartment association of the contacting domains into A-A, A-B and B-B. Contacts within domains less than 2Mb apart were discarded. We summarized the statistics of observed total compartment contact per cell as (O_AA_, O_AB_, O_BB_). The compartment score of a cell is the depletion of A-B contacts over the count expected by the marginal distribution of A or B contacts:

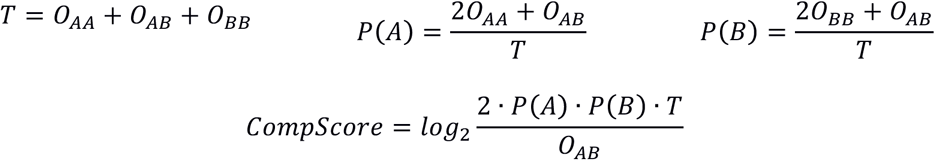

### Analyzing CTCF loops

We followed these steps to generate a set of loops:

- Genomic regions with strong CTCF ChIP-seq^16^ signal were screened (regions on which either of the two ChIP-seq replicates quantile is larger than 99%)
- Strong CTCF motifs were screened (Motif strength > 99.988% of the genome-wide distribution)
- Each 50bp window was classified as *C* (only ChIP), *F* (ChIP + forward motif), *R* (ChIP + reverse motif) or *B* (ChIP + forward and reverse motif).
- Each ChIP-supported motif is classified by its upstream and downstream (200Kb) mean time of replication (ToR) as early (if ToR is positive) or late (if ToR is negative), resulting in the following anchor counts (discarding 116 loops with missing ToR information):
- Convergently oriented ChIP-supported motifs (B/F anchor upstream to a B/R one) distanced 200Kb-1Mb from each other were considered as loop candidates
- Loop candidates were further filtered by coverage enrichment in the normalized pooled contact map, requiring contact enrichment score higher than 60 within a 20×20Kb window centered on the loop. This resulted in a set of 53361 loops, with the following anchors classification:

| Upstream ToR | Downstream ToR | Denoted | Count |
| --- | --- | --- | --- |
| Early | Early | EE | 55,778 |
| Early | Late | EL | 2,459 |
| Late | Early | LE | 2,598 |
| Late | Late | LL | 10,278 |

| 1<sup>st</sup> anchor | 2<sup>nd</sup> anchor | Denoted | Count |
| --- | --- | --- | --- |
| EE | EE | Early-Early | 37,914 |
| EE | LL | Early-Late | 2,295 |
| LL | EE | Late-Early | 2,187 |
| LL | LL | Late-Late | 3,191 |
| Mixed (EL/LE) | Mixed (EL/LE) | Mixed | 8,135 |

Loop foci enrichment (**Fig. 3d**) quantifies how concentrated contacts are around a loop. We calculate the ratio between contacts in a small window (20×20kb) centered on the loop and contacts in a larger window (60×60kb), normalizing by the expected ratio if contacts were uniformly distributed (1/9). To get a mean loop foci enrichment over a group of loops the sum of contacts in all small windows is compared with the sum of contacts in the larger ones.

Spatial analysis of contact distribution around loops (**Fig. 3e**) was performed by grouping observed and shuffled contacts in a 50×50kb window around loops anchors into 2Kb-14Kb square bins and color coding according to the per-bin log fold ratios over shuffled-data statistics.

### Domain condensation

The condensation level of a domain is quantified by the mean contact distance of contacts within the domain. To ensure domain coherence for condensation analysis (**Fig. 3f**) we used a more fine-grained domain definition with a higher threshold on the insulation (85 percentile for calling domains, instead of 90) resulting in 3502 domains, out of which 3434 had time of replication information. Each domain was classified by its mean time of replication and its length, resulting in the following groups of domains:

|  |  | Domains length range |  |  |  |
| --- | --- | --- | --- | --- | --- |
|  | mean ToR (mT) | 1-244K | 244-490K | 490-980K | 980-1500K |
| **Early** | mT > 0.84 | 346 | 401 | 229 | 29 |
| **Mid-Early** | 0 < mT < 0.84 | 336 | 329 | 270 | 52 |
| **Mid-Late** | -0.66 < mT < 0 | 146 | 166 | 213 | 104 |
| **Late** | mT < -0.66 | 115 | 124 | 198 | 109 |
| **Total** |  | 943 | 1020 | 910 | 294 |

As expected from earlier reports, domains length was correlated with time of replication, but sufficient statistics was available for analysis in less populated bins:

|  | **1-244K** | **244-490K** | **490-980K** | **980-1500K** |
| --- | --- | --- | --- | --- |
| **Early** | 36.7% | 39.3% | 25.2% | 9.9% |
| **Mid-Early** | 35.6% | 32.3% | 29.7% | 17.7% |
| **Mid-Late** | 15.5% | 16.3% | 23.4% | 35.4% |
| **Late** | 12.2% | 12.2% | 21.8% | 37.1% |

### Computing distributions of single-cell compartment association scores

In order to analyze the compartmentalization behavior of TADs in a population of single cells, we initially classified each TAD to the A or B compartment according to its time of replication, (A - Repli-chip > 0). Next, for each TAD we randomly selected 40 cells in which at least 6 contacts associated the TAD intra-chromosomally with TADs at least 2Mb away. A given TAD‘s compartment association score for a given cell is the number of contacts (out of the six sampled) that represent an interaction with a TAD labeled A in the initial binary compartment assignment. For example, a TAD that only interacted with B TADs receives a score of zero and one that interacted solely with A TADs receives a score of six, The distribution of association scores across the sampled cells (different cells per TAD) represents the overall tendency of the TAD to be strongly associated with A, B. This procedure was repeated using cells categorized by our phasing as G1, early to mid S, and late S to G2, producing for each TAD a distribution of compartment association scores across the three cell cycle phases. We note that down-sampling (to a fixed number of cells and a fixed number of long-range intra-chromosomal contacts per cell) is important for controlling for coverage biases. The parameters here (40 and 6) were selected to ensure a sufficient number of TADs are represented by the analysis. Overall, 2649 TADs were analyzed in the haploid dataset.

### Ranking TADs by compartment association score distributions and assigning to pseudo-compartments

For unsupervised analysis, TADs were clustered by the distribution of their compartment association scores in each group of cells (**Extended Data Fig. 9a**). Given the absence of clusters showing bimodal distributions, TADs were ranked by the difference between the summed values of the two most-A bins and the summed values of the two most-B bins in the late S-G2 distribution, with the analogous difference in the early S and G1 distributions serving as tiebreakers where needed. Pseudo-compartments were then defined by discretizing the ranking into 20 groups. We assigned TADs that were not included in the original downsampled analysis to pseudo-compartments by majority voting on their *cis* contacts in the ensemble map.

### Creating normalized pseudo-compartment interaction maps

We derived an observed pseudo-compartment interaction map from the observed ensemble map for each cell-cycle phase by summing the number of contacts between domains in each pseudo-compartment, separately for *cis*- and for *trans*-chromosomal contacts. To derive the *cis* pseudo-compartment maps in **Fig. 4e**, we took the log of the element-wise ratio between the observed pseudo-compartment map and an expected pseudo-compartment map (summed from the shuffled pool map). For *trans*-chromosomal contacts we used the expected contingency table from the observed pseudo-compartment *trans* contact map, taking the log ratio of observed and expected to obtain the *trans* pseudo-compartment enrichment maps shown in **Fig. 4f**.

### Epigenetic annotations

| Track | Source | Usage | Processing |
| --- | --- | --- | --- |
| Time of replication | Repli-chip of ES-D3<sup>16</sup> (rep 1) | Calculating repli-score and classifying TADs | Mean value on regions (TADs, upstream/downstream to landmarks, 10kb around a fend for repli-score) |
| Haploid RNA-seq | In-house (experimental details above) | 4g | Log2(mean of 2 replicates over TAD) |
| Diploid RNA-seq | In-house (experimental details above) | Ext 3d | -log2(1 – max(reps max percentile) |
| H3K4me3 | ChIP-seq of ES-Bruce4<sup>16</sup> | 4g | Mean of 2 replicates over TAD |
| H3K4me3 | ChIP-seq of ES-Bruce4<sup>16</sup> (rep 1) | Ext 3d | Fraction of TAD with peaks (peak = score > 99% percentile) |
| H3K27me3 | ChIP-seq of ES-Bruce4<sup>16</sup> | 4g, Ext 3d | Mean of 2 replicates over TAD |
| LaminB1 | mESC LaminB1 DamID<sup>24</sup> | 4g, Ext 3d | mean LaminB1 value over TAD |
| CTCF | ChIP-seq of ES-Bruce4<sup>16</sup> | Fig 3d-e | Details on loops section above |

### Mapping and filtering before 3D modelling

Since structure modeling is sensitive to low levels of noise, haploid G1 cells were processed using stringent contact filtering to remove contacts that are more likely to be technical artifacts. We first used HiCUP^25^, applying a di-tag size selection from 50 bps to 850bps, for mapping di-tags and filtering out common Hi-C artefacts. Putative PCR duplicates were not removed by HiCUP, instead the filtered data was then passed a new tool (SiCUP) for further single-cell Hi-C specific filtering. We removed reads mapping to the Y chromosome, to short restriction fragment (less than 21bps) and to regions defined as problematic by ENCODE. We also filtered reads mapping to fragment ends forming multiple interactions in one percent or more of the datasets. To avoid potential artefacts we removed singleton di-tags. In haploid G1 cells there is only one copy of the genome, hence after removal of PCR-duplicates each observed fragment end should be in contact with at most one other fragment end. Consequently, multiple contacts from the same fragment were removed entirely. An exception to this was when a fragment end (A) interacted with two other fragments ends (B and C) which were close together (defined here as when B and C were within 20 MboI fragments). In such instances the strand orientation of the reads mapping to B and C were typically the same, to a degree not expected by chance (as defined by a chi-squared test when evaluating the whole dataset). We reasoned that in such instances these apparently distinct interactions were in fact derived from one initial Hi-C interaction. Consequently, when this was observed, not all the di-tags were discarded. Instead, if the Hi-C interaction was in trans, a random di-tag was discarded. Alternatively, when the Hi-C interaction was in cis, the di-tag representing the shortest Hi-C interaction was retained.

We also filtered out unsupported contacts. For each cell, using the filtered contacts, we first derived a connectivity graph of the genome. Nodes of the graph represented 1Mb segments of the genome, and each edge represented a single contact mapped onto 1 Mb resolution, so any two nodes of the graph might be connected by more than one edge. We defined a contact unsupported if upon deletion of that contact the shortest path connecting its two end nodes would be longer than 3 edges. These unsupported contacts (median 1.06% of contacts, **Extended Data Fig. 10a**) were removed from the sc-HiC libraries before 3D modelling.

### Polymer model for whole-genome modelling

We modeled chromatin as a simple non-overlapping beads-on-a-string polymer^11^ subject to data derived constraints. The energy function consists of a bond potential that ensures the connectivity of the polymer chain within chromosomes, a second term that prevents overlaps of chromatin and a third term that enforces the observed contacts as distance constraints with different contact points for each cell. We implemented this coarse-grained polymer model to be used with the GROMACS 5.0.6 molecular dynamics package^26^. The energy function and the exact values of the model parameters are listed in the Supplementary Material. Briefly, the exclusion energy term is a harmonic function for distances below *d_0_* with a force constant *k*^excl^, and the bond and constraint energy terms are defined by a flat-bottomed harmonic function with no energy penalty for distances between *d*_0_ to *d*_1_, and a harmonic potential with a force constant *k* outside this range which turns into a linear function for distances greater than *d*_2_. 3D models were optimised using a simulated annealing protocol, using a piecewise linear temperature profile, and different initial conditions to assess convergence robustness.

### Quality control of 3D models

To avoid underdetermined models, we only considered models that were computed using at least 10,000 filtered contacts. The root mean square distance (RMSD, the smallest possible Euclidean distance) of models, indicating the reproducibility of models, as a function of the number of contacts is shown in **External Data Fig. 10d**. We also discarded models with high numbers of violated constraints. For this, we considered a constraint violated when its distance was greater than *d*_2_ (constrained distances are expected to be greater than *d*_1_ by construction, due to thermal fluctuations). At 1 Mb resolution there were no violated constraints for 171 of the 190 cells (**External Data Fig. 10b**), and the fraction of violated constraints strongly correlated at the 500 kb and 100 kb resolutions (Pearson R = 0.96). For further analysis, we selected 127 cells that on median had no more than 0.5% violated constraints at the 500 kb resolution and no more than 0.1% violated constraints at the 100 kb resolution (shown as black dots in **External Data Fig. 10c**). We further discarded one cell that had more than 10% of a chromosome missing, resulting in 126 ordered G1 cells with high-quality models for downstream analysis.

We tested robustness to the initial conditions by using permuted chromosomes along a spiral as well as folded structures of other nuclei as the initial chromatin conformation. Results were robust to changes to the initial conditions, indicating the convergence of the simulations, which was measured by computing the distribution of the root-mean-square-distance (RMSD) between aligned models of the same cell compared to different cells (**External Data Fig. 10e**). Models at different resolutions were compared to one another by down-sampling the higher resolution structures to 1 Mb, and scaling the nuclear radius of the model to the nuclear radius of the 1 Mb model. The down-sampled models computed at different resolutions were consistent at 1 Mb resolution. Models were also robust when scaling the model parameters by a factor of half or two.

### Cross validation

To test the robustness of our 3D modelling approach to missing contacts, we performed a cross validation test. For each cell, we identified the 5 most strongly interacting chromosome pairs, measured by the number of contacts between the two chromosomes. For each of these 5 chromosome pairs, we removed all *trans* contacts between that chromosome pair and recomputed a structure model for that cell (we only delete interactions between one chromosome pair at a time). We computed the distance distribution of the removed constraints in the models computed with those constraints removed (**External Data Fig. 10f** blue line), and compared it to their distance distribution in randomly selected cells (control, **External Data Fig. 10f** green line). The distances of these removed contacts are noticeably shorter in their corresponding model than they are in randomly selected other cell, indicating that the proximity of the chromosome pair is preserved. This suggests that these sparse datasets contain sufficient information about chromosome level interactions for robust structural modelling. This is probably due to their indirect coupling through their contacts with their neighbouring chromosomes. In comparison, the distances of the unsupported contacts filtered out before modelling (**External Data Fig. 10f** light blue line) were much more similar to the control, indicating that most of those interactions are likely noise.

### Chromosome decondensation

We grouped the 126 cells into 7 time groups in G1 (**External Data Fig. 11a**). As the chromatin structure varied most in early G1, we used 4 time groups in the first quarter of the 126 G1 ordered cells, and 1 group for each of the remaining three quarters. We computed the nuclear radius from its radius of gyration. As an indicator of nucleus sphericality, we also computed the inertia ellipsoid of the nucleus. The detailed description of how these quantities are computed can be found in the Supplementary Material. We noted that although the nuclear lamina is not modelled by the force field, the models turn out to be near spherical (a ≈ b ≈ c) and roughly the same size (**External Data Fig. 10g**, nucleus a/c ratio). We also noted that low mappability regions tend to be more flexible and uncertain in their position in the computed models, illustrated by the unconstrained region around chrX:30Mb (the white chromatin region looping out of the models in **External Data Fig. 10e)**. To illustrate changes in the overall chromatin structure, we selected an M-phase and a late (118^th^ of the 126 ordered cells) G1 cell. We plotted the whole genome structure and a karyotyped image at 100 kb resolution (**Fig. 5a**). The structure models were visualised and aligned using VMD^27^ 1.9.1, the images were rendered using PovRay 3.7 (http://www.povray.org). The colour scheme used for the chromosomes was chosen to provide maximum contrast^28^.

To quantify the decondensation of chromosomes during G1, we fitted an ellipsoid to each chromosome of every model and averaged the values over all models of the same cell. As a measure of sphericality, we used the longest-to-shortest semiaxis ratio (**External Data Fig. 10g**, chromosome a/c ratio) together with the middle-to- shortest semiaxis ratio (**External Data Fig. 10g**, chromosome b/c ratio). Larger a/c ratios together with b/c ≈ 1 in early G1 indicate rod-like structures that decondense into a more spherical structure as a/c decreases to a ≈ b ≈ c. We plotted the distribution of all chromosome a/c ratios over cells in the 7 time groups with illustrating examples of chromosome 1 from each time group in **Fig. 5b** of the Main text.

### 3D organisation of compartments

To investigate the spatial organisation of compartmentalisation, we assigned compartment labels to the beads using a majority vote. Beads with no compartments were excluded from further analysis. As an indicator of local decompaction of chromatin, we computed the average distance of neighbouring chromatin segments of the same compartment, normalised by the d_1_^bond^ bond length parameter. We computed the average compartment decompaction values for each model, and averaged them across all 3D models of the same cell. The distribution over all cells in the 7 time groups is shown in **Fig. 5c,d**. We tested if the mean A and B compartment decompaction values came from the same distribution using Kolmogorov-Smirnov statistics.

We also computed the average radial positioning of the compartments. For each bead, we computed its radial position as a volume fraction, increasing linearly between the nucleus centre-of-mass (0) and the nuclear radius (1) in the volume enclosed by a sphere of that distance radius from the nucleus centre. We averaged the radial position values of all beads of the same compartment, and across all 3D models of the same cell. We plotted the distribution of the radial positions across the cells in the 7 time groups in **Fig. 5e,f**. We tested if the mean A and B compartment radial position values came from the same distribution.

Testing the enrichment of compartment co-localisation between pairs of pseudo-compartments in long *cis* (> 2 Mb genomic distance) or *trans* was done separately. For this analysis, we defined two beads in contact if their distance was within 10% of the nucleus diameter. For any two pseudo-compartments, we counted the contacting bead pairs in long *cis* or *trans,* for which one bead belonged to one pseudo-compartment, while the other bead belonged to the other pseudo-compartment. We averaged these counts over all models of all cells in the 7 time groups of G1, and rounded them down to the nearest integer, resulting the observed counts. We shuffled the bead contacts while keeping the number of contact beads of each compartment constant, resulting the expected counts. We plotted the log_2_(observed/expected) ratios between any two pseudo-compartments as a matrix (**Fig. 5g,h**).

### Nuclear Organiser Region clustering

For each chromosome, we used the first constrained bead as the pericentromeric region closest to the centromere. Chromosomes were divided into two groups, Nucleolar Organiser Region (NOR) chromosomes (chromosomes 11, 12, 15, 16, 18 and 19), and other chromosomes^17^. We computed all pairwise distances between the pericentromeric regions within these two groups normalised by the nucleus diameter, and plotted the distribution of the top one third of these distances across all models of cells for each time group (**External Data Fig. 11b**). We averaged these distances for each model and each cell, and tested if the mean of the top third shortest centromere distances came from the same distribution by performing a Kolmogorov-Smirnov test on each time group.

**Extended Data Fig. 1:**
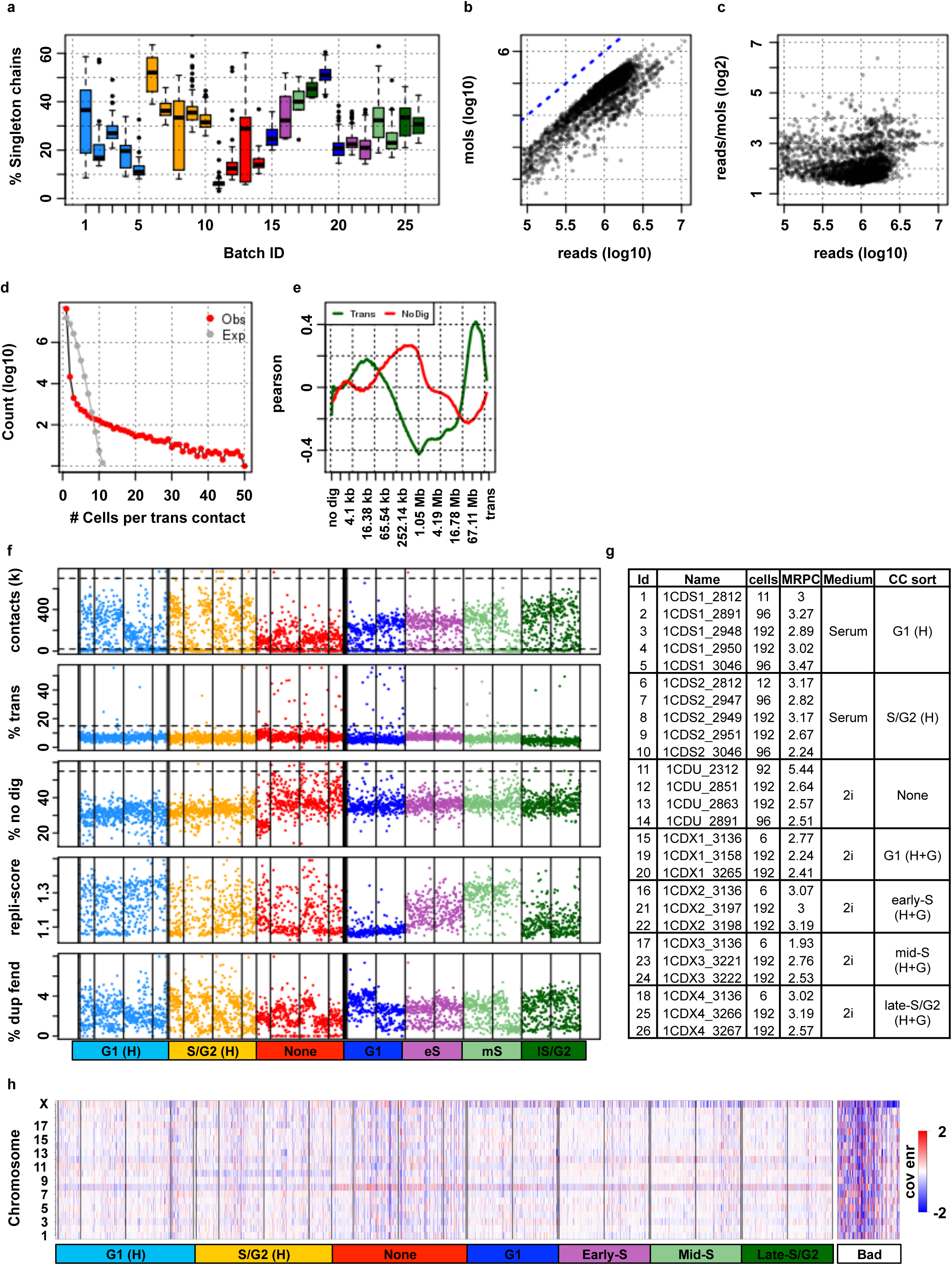
Technical QCs. **a)** Testing library saturation. Showing the fraction of segment chains supported by a single read per batch, batch colors match their sorting criteria. **b)** Number of unique molecules captured per cell against the number of sequenced reads. **c)** Number of reads per unique molecule against number of sequenced reads. **d)** Observed (red) and expected (by binomial reshuffling of the observed contacts) number of cells each trans contact appears in. **e)** Correlation of the fraction of trans contacts and close ***cis*** ( < 1 kb, non-digested) with the fraction of contacts in different distance bins. **f)** QC metrics of single cells colored by FACS sorting criteria: G1 (H) and S/G2 (orange) by Hoechst staining, G1, early-S (eS), mid-S (mS) and late-S/G2 (lS/ G2) by Hoechst and Geminin staining (see color legend at bottom). Vertical lines mark experimental batches. Showing from top to bottom total number of contacts (coverage); fraction of inter-chromosomal contacts ***(%trans);*** fraction of very close (< 1 Kb) intrachromosomal contacts (%no dig); Early replicating coverage enrichment (repli-score); fraction of fragment ends (fends) covered more than once (%dup fend). Horizontal dashed lines mark thresholds used to Alter good cells. **g)** Details on the experimental batches, showing the number of cells in each batch, mean number of reads per cell (MRPC, in million reads), medium used to grow the cells and FACS sorting criteria (H = Hoechst, G = Geminin). **h)** Coverage enrichment per cell (column) and chromosome (rows), expected coverage is calculated from the frequency in the pooled cells and the total number of contacts in each cell. We discard cells that have any aberrantly covered chromosome (at least 2 fold enrichment or depletion, shown on the right “Bad” panel), left panel shows all cells.

**Extended Data Fig. 2:**
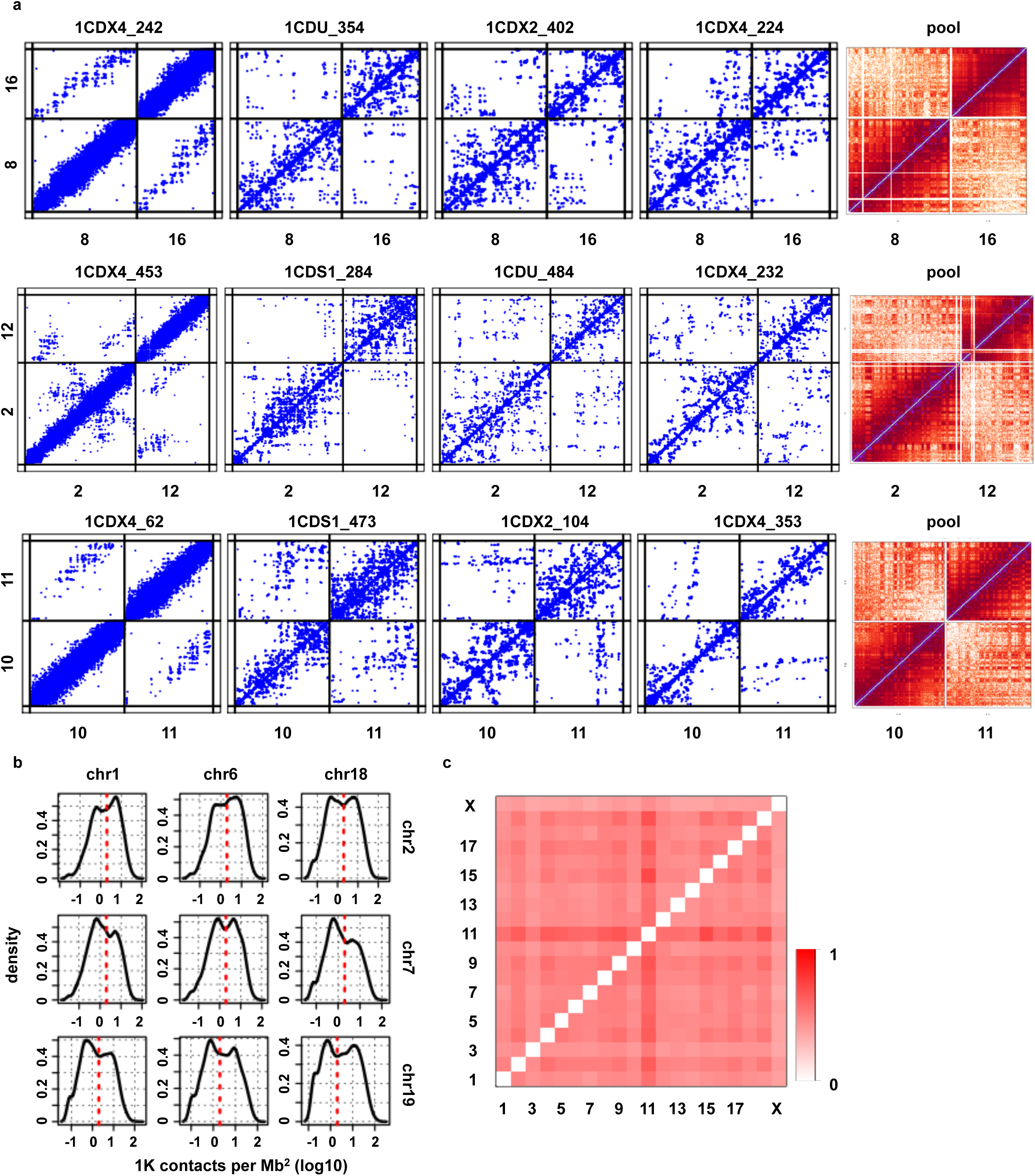
Trans-chromosomal contacts. **a)** Examples of inter-chromosomal contact map for several single cells (blue) and the pooled contact map on the same chromosomes. **b)** Distribution of number of contacts between selected pairs of chromosomes. Showing the number of contacts per square Mb, red dashed line marks the cutoff for interacting chromosomes. **c)** Fraction of cells in which each pair of chromosomes was interacting (had normalised interaction above the cutoff shown in panel **b)**.

**Extended Data Fig. 3:**
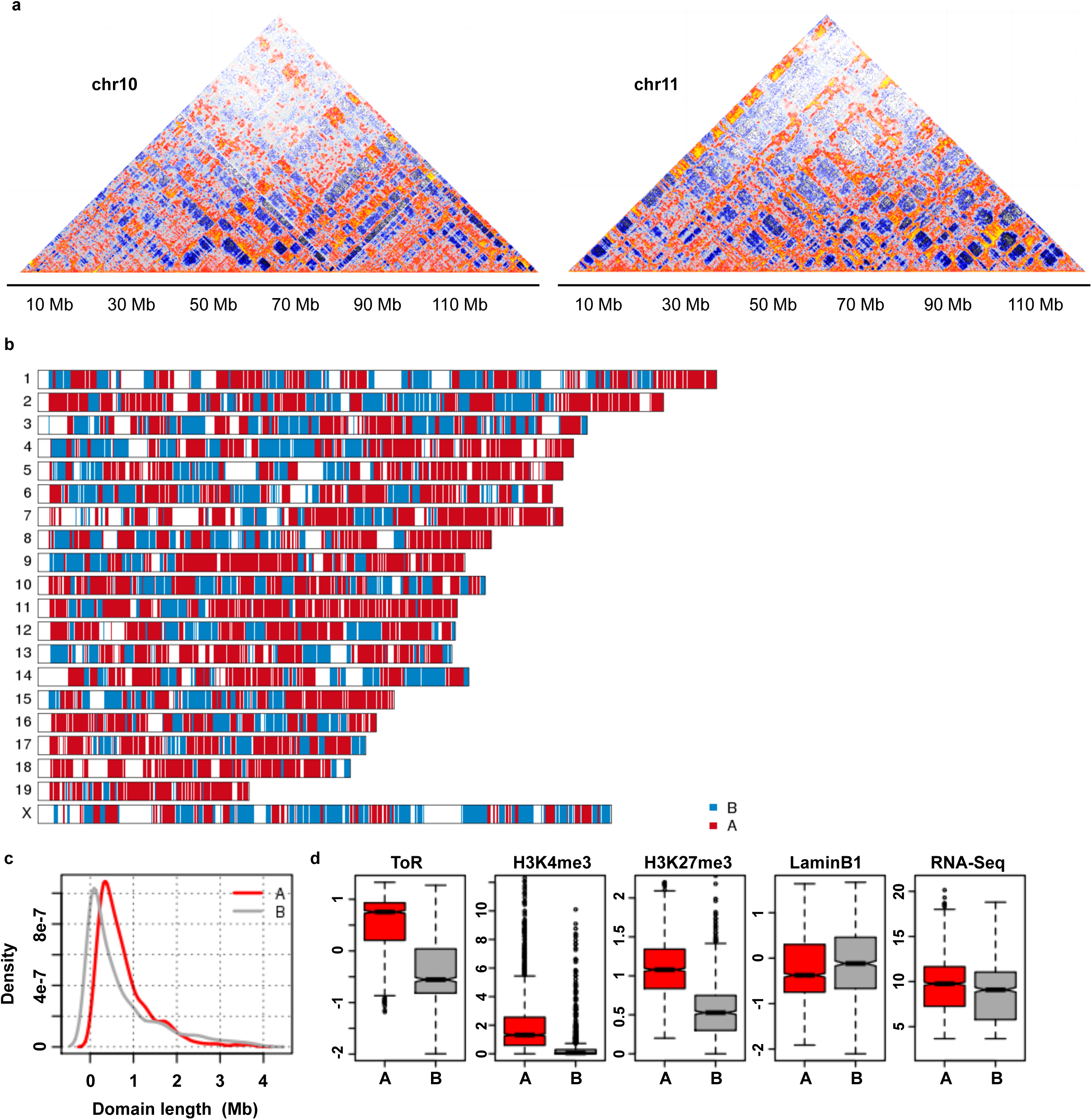
Pooled contact map and population data. **a)** Normalised chromosomal contact maps of the pooled single cells (showing chromosomes 10 and 11). **b)** Ideogram showing the division of domains to the A and B compartments by k- means clustering of domains irans-chromosomal contact profiles, **c)** Comparing A and B domains by their lengths, **d)** Comparing A and B domains by their mean time of replication (ToR); percentage of H3K4me3 peaks; mean H3K27me3; mean LaminBI values; mean RNA-Seq.

**Extended Data Fig. 4:**
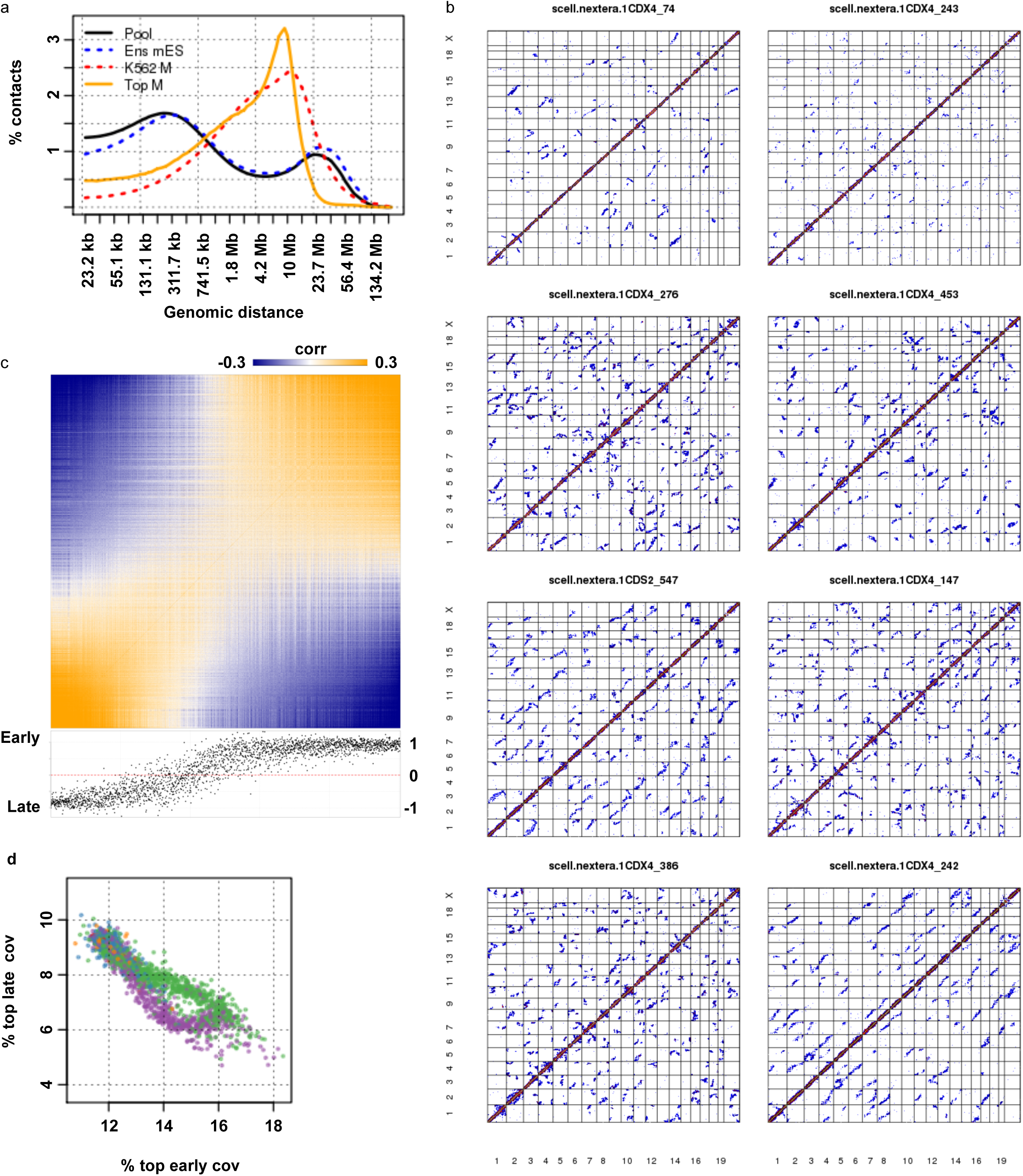
Mitotic cells. **a)** Contact decay profile of our pool of singles (Pool), a reference mouse ESC ensemble HiC dataset (Ens mES), Naumova et al. (reference 12) human K562 mitotic Hi-C (K562 M) and our pooled 8 most mitotic-like cells (Top M). **b)** Genome-wide contact maps of 8 putative mitotic diploid cells (cells with the highest fraction of contacts at 2-12 Mb distances). **c)** Correlation matrix of domain coverage across cells. Ordered by the mean correlation to the top 50 earliest replicating domains minus the mean correlation with the latest 50 domains. Time of replication of each domain is shown at the bottom. **d)** Comparing the fraction of contacts associated with the latest- and earliest- replicating fends (ToR < -0.5 and ToR > 0.5, respectively), cells are colored as in **Fig. 1e**.

**Extended Data Fig. 5:**
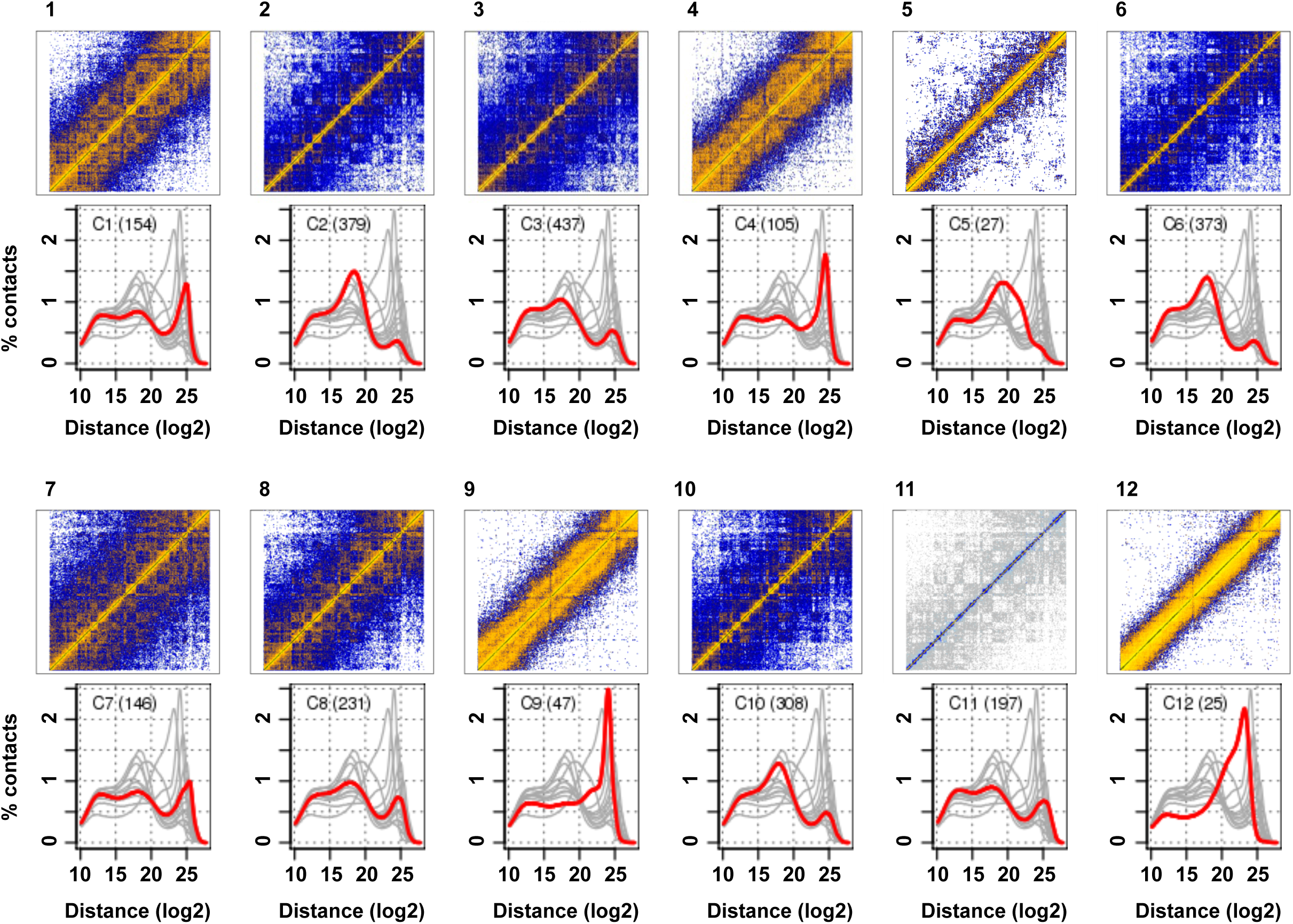
Clustering cells by contact decay. Chromosomal contact maps (chromosome 6) of pooled cells per cluster and the decay trend of each cluster (red) compared to the rest of clusters (grey).

**Extended Data Fig. 6:**
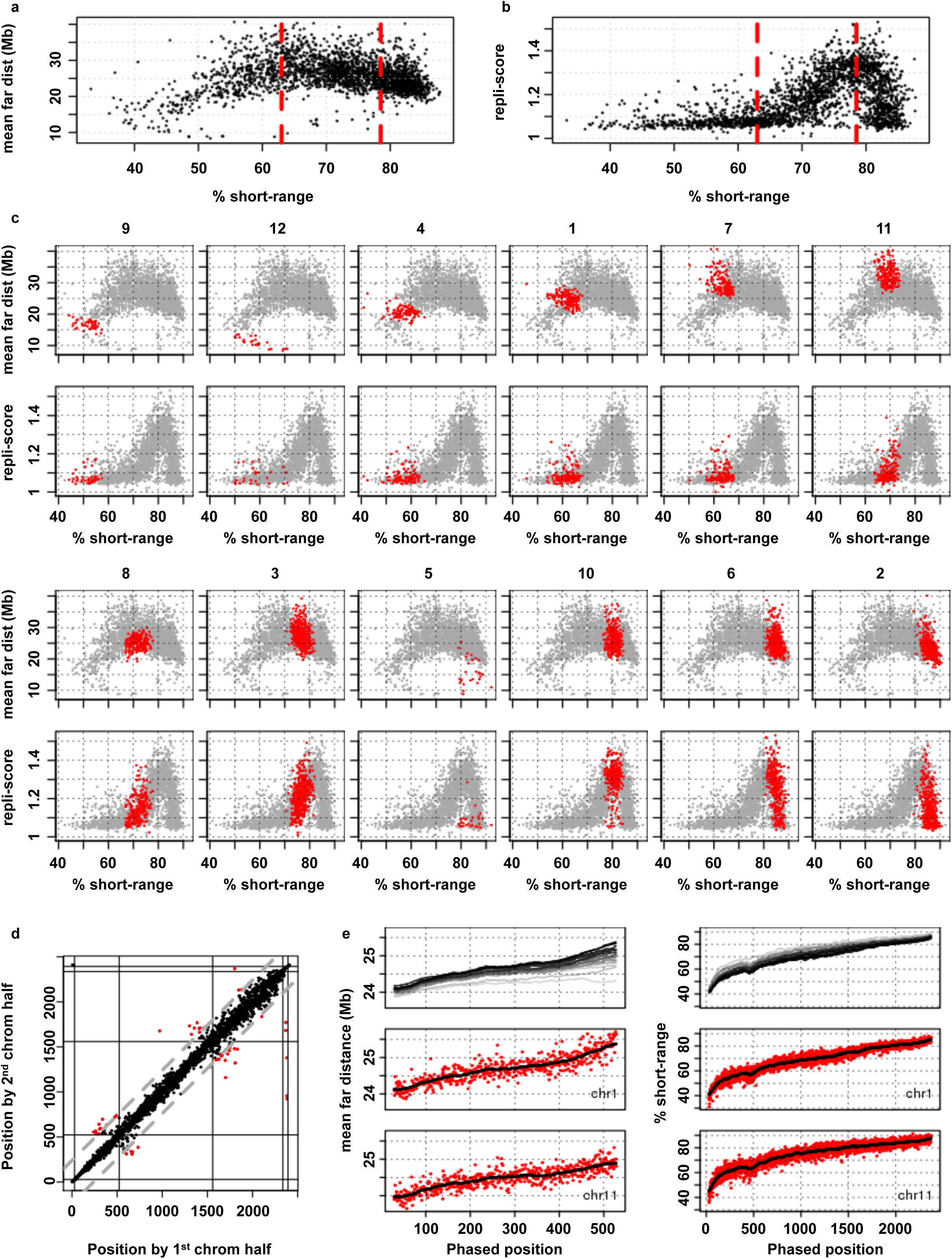
Phasing cells over the cell cycle. **a)** Mean contact distance among the far (4.5-225 Mb) range against the fraction of short-range (32 kb-2 Mb) contacts, dashed red lines mark the cutoffs used to divide cells into the main 3 groups (G1, early-S, mid-S/G2). **b)** Similar to panel **a)** but showing repli-score against the fraction of short-range contacts. **c)** Mean far contact distances and repli-score against the fraction of short-range contacts per Extended Data 5 clusters (red - cells in cluster, grey - all cells), ordering clusters by their mean %short-range. **d)** Testing the stability of our phasing, we compare the positions of cells by phasing using only half the chromosomes (1^st^ half: 2, 3, 5, 6, 9, 10, 12, 15, 17, 19, 2^nd^ half: X, 1, 4, 7, 8, 11, 13, 14, 16, 18). Position of only 54 cells differ in more than 10% of the number of cells (outliers marked in red, 10% margin in dashed grey). Phasing groups marked in black lines. **e)** Chromosomal comparison of the contact decay metrics used to phase the cells in each group: mean far contact distance for the G1 cells (left) and %short-range for G1, S and G2 cells (right). Showing smoothed (n = 51) trends per chromosome on top, chromosomes colored by length (light grey - chromosome 19, black - chromosome 1). Data for specific chromosomes: chromosome 1 (middle) and chromosome 11 (bottom) as red dots.

**Extended Data Fig. 7:**
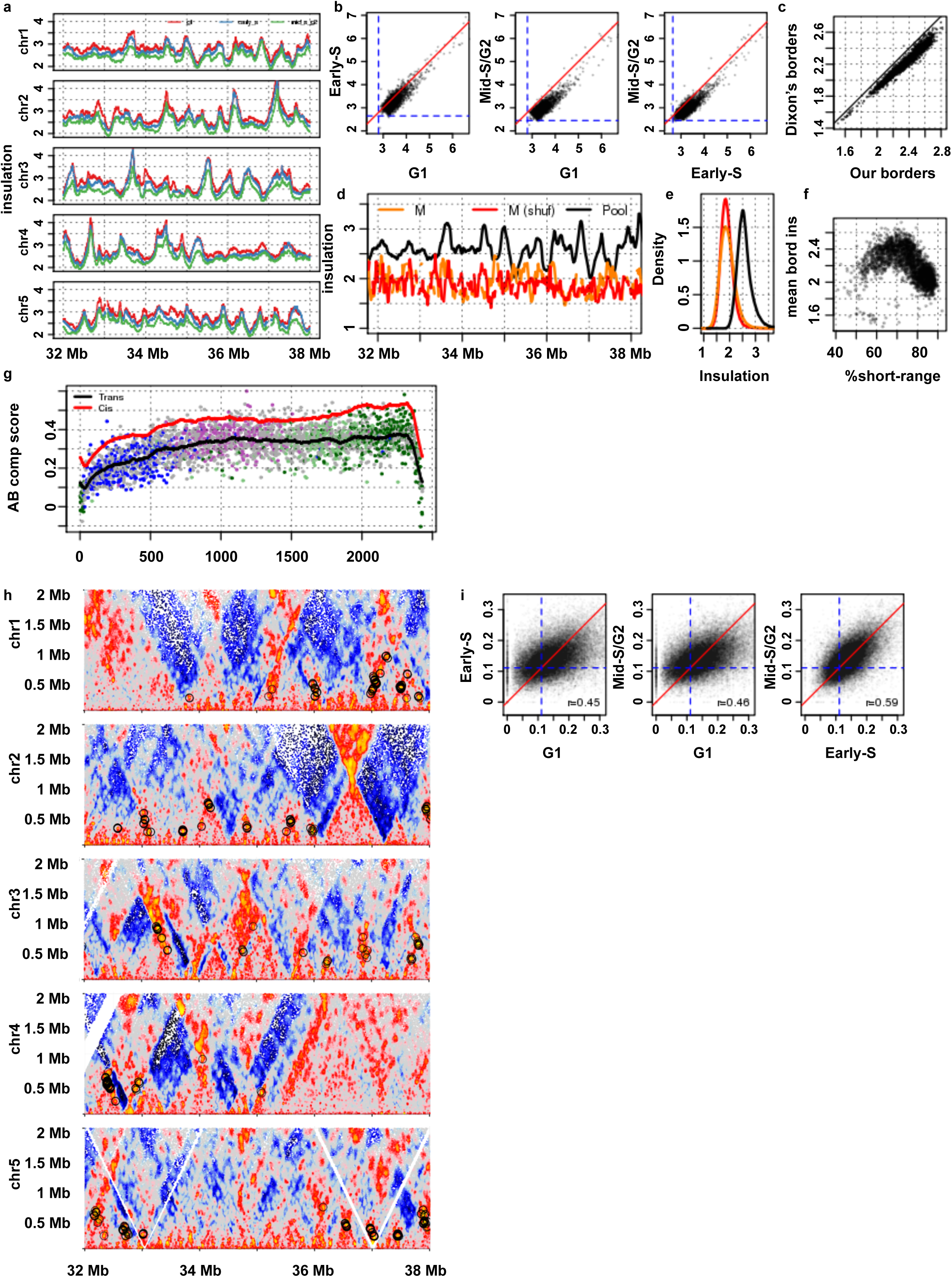
Insulation of sub-groups and loops. **a)** Insulation profiles of the large phased groups over 6 Mb regions in chromosomes 1-5. **b)** Comparing border insulation of phased groups. Dashed blue lines show the mean insulation value per group. **c)** Showing mean insulation per cell on mouse ESC TAD borders taken from Dixon ***et al.*** 2013 (reference 5; using the center point of borders smaller than 80 kb) compared to the mean cell insulation over our inferred borders. **d)** Insulation profile over a 6 Mb region in chromosome 1 (same as panel a), showing insulation of pooled cells (black), pooled mitotic cells (red) and a shuffled pool of mitotic cells (orange). **e)** Genome-wide distribution of insulation values for pooled-, pooled-mitotic- and shuffled- pooled-mitotic cells, same colors as in panel **c**. **f)** Mean border insulation per cell against the fraction of short-range (< 2 Mb) contacts. **g)** A/B compartment score on trans-chromosomal contacts, single cell mean values in dots colored by FACS sorting, mean trend (black) and mean trend of c/s-chromosomal A/B compartment in red (same trend as in **Fig. 3a**). **h)** Normalized contact maps of the regions as in panel **a**, circling the loops detected in distances 200 kb – 1 Mb. **i)** Comparing loop foci enrichment calculated per phased group, non-enriched value marked in dashed blue.

**Extended Data Fig. 8:**
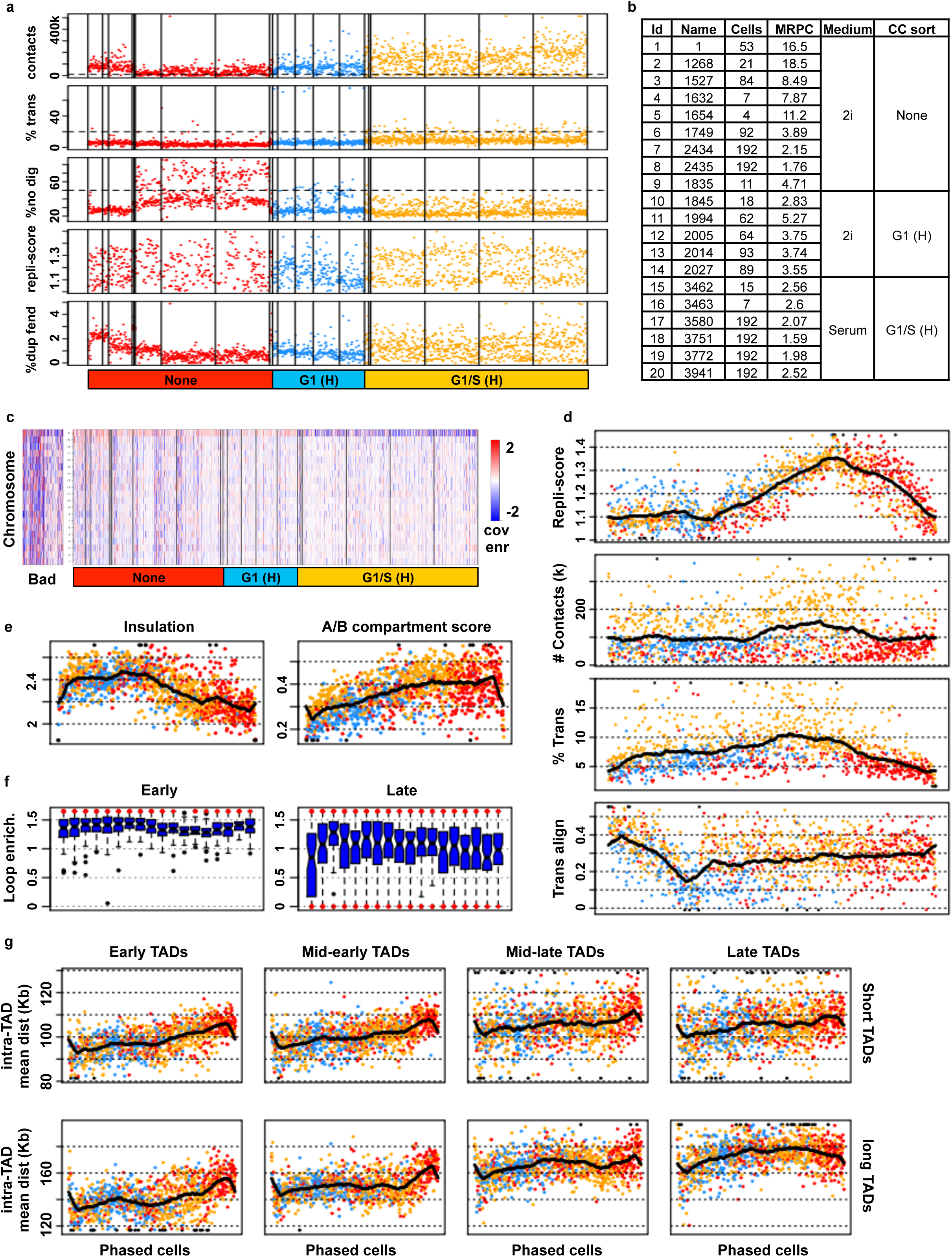
Haploid cells technical QCs. **a)** QC metrics of single cells colored by FACS sorting criteria: G1 (H) and G1/S (H) by Hoechst staining, (see color legend at bottom). Vertical lines mark experimental batches. Showing from top to bottom total number of contacts (coverage); fraction of inter-chromosomal contacts ***(%trans);*** fraction of very close (< 1 kb) intrachromosomal contacts (%no dig); Early replicating coverage enrichment (repli-score); fraction of fragment ends (fends) covered more than once (%dup fend). Horizontal dashed lines mark thresholds used to filter good cells. **b)** Details on the experimental batches, showing the number of cells in each batch, mean number of reads per cell (MRPC, in million reads), medium used to grow the cells and FACS sorting criteria (H = Hoechst). **c)** Coverage enrichment per cell (column) and chromosome (rows), expected coverage is calculated from the pooled cells. We discard cells that have any aberrantly covered chromosome (at least 2 fold enrichment or depletion, shown on the left “Bad” panel), right panel shows all cells. **d)** Similar to **Fig. 2c** but produced from the haploid cells, outliers colored in black. **e)** Similar to **Fig. 3a** but produced from the haploid cells, outliers colored in black. **f)** Similar to ****Fig. 3d**** but produced from the haploid cells. **g)** Similar to **Fig. 3f** but produced from the haploid cells, outliers colored in black.

**Extended Data Fig. 9:**
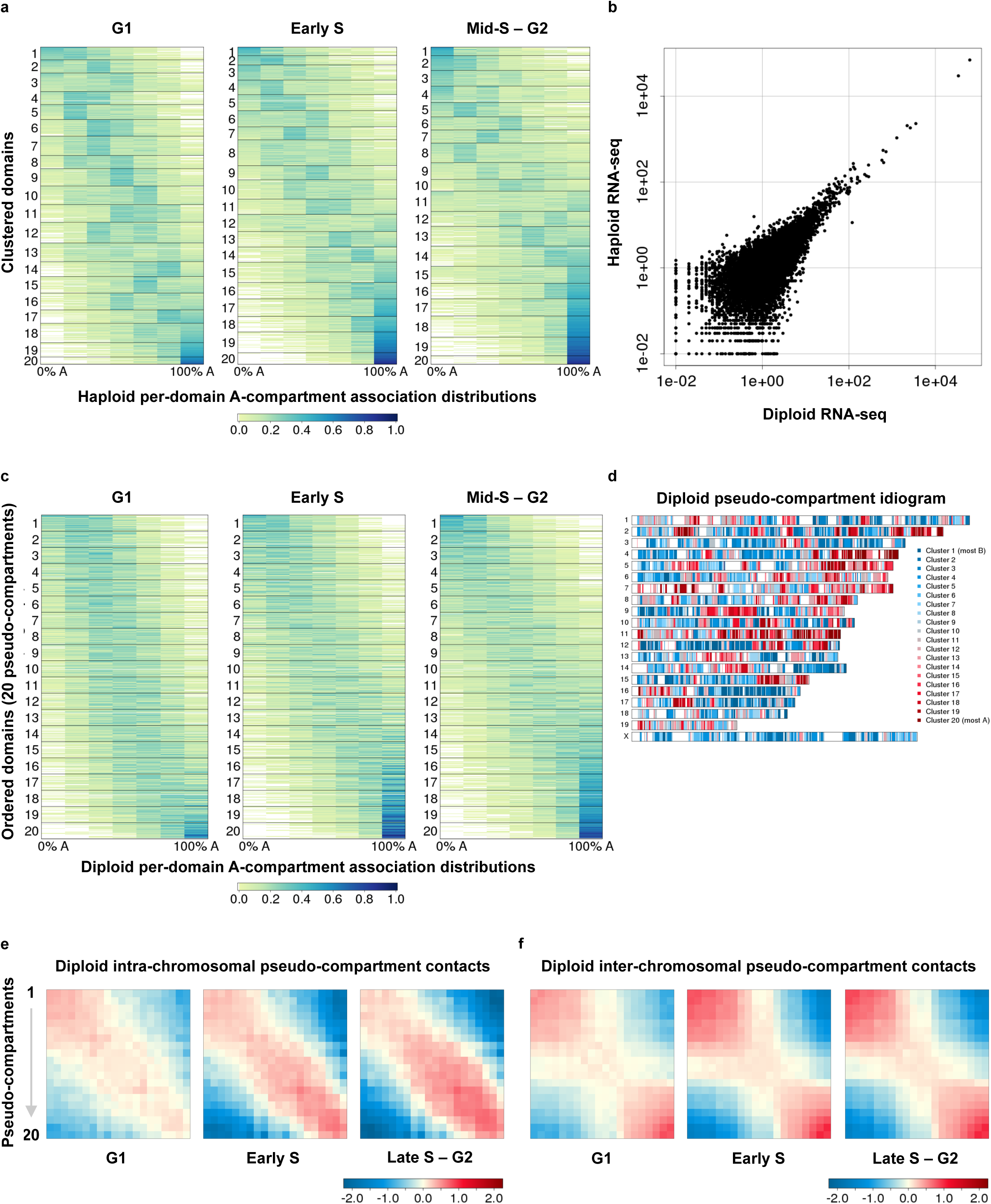
Haploid cells clustering, diploid cells pseudo-compartment analysis. **a)** Per-domain distributions of A-compartment association across haploid cells in each of the three FAC-sorted cell populations (from left to right: G1, early S, late S to G2). For each TAD we sampled 6 long-range contacts (over 2 Mb) in each nucleus, and counted the number of A-compartment associations out of these. The distributions of a given domain' s percentage of A-associated contacts over the sampled cells in the three cell-cycle phase groups are shown as a row in the heatmap. Distributions were clustered using fc-means (performed separately for each cell cycle phase) to 20 clusters. **b)** Haploid vs. diploid RNA-seq data. **c)** Per-domain distributions of A- compartment association across diploid cells in each of the three FAC-sorted cell populations (from left to right: G1, early S, late S to G2). Domains were ordered on the basis of their distribution in late S/G2, and the ordered domains then divided into 20 pseudocompartment groups. **d)** Pseudo-compartment labeling of all diploid domains. **e)** Normalized (log enrichment) pooled diploid contact maps of intra-chromosomal interactions between domains in each pseudo-compartment. **f)** Normalized (log enrichment) pooled diploid contact maps of irans-chromosomal interactions between domains in each pseudo-compartment.

**Extended Data Fig. 10:**
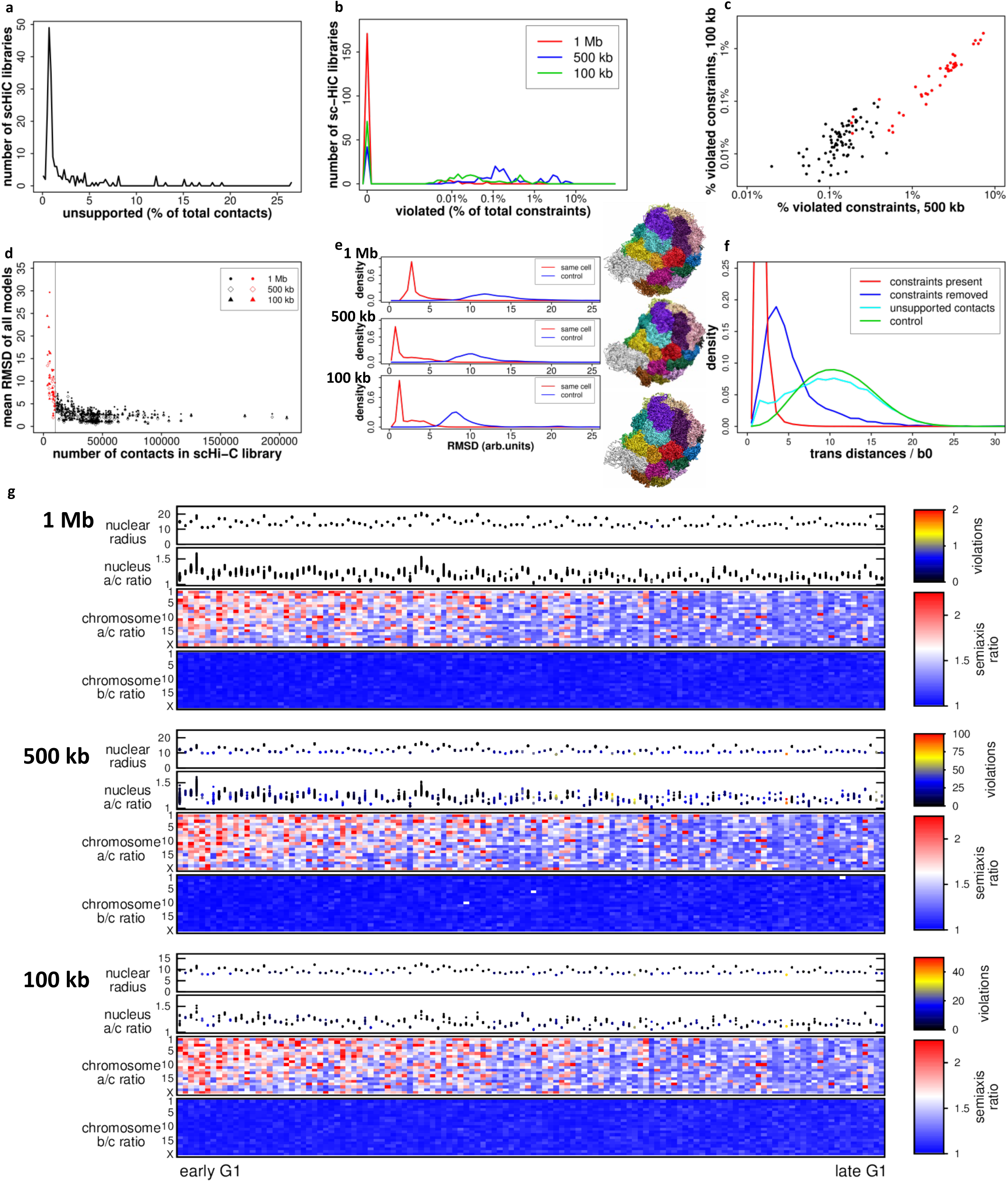
Quality control for whole genome structure modeling. **a)** The distribution of the fraction of unsupported contacts in the pre-filtered contact set in all 190 haploid cells in 2i inferred as being in G1. **b)** The distribution of the fraction of violated constraints in the 190 haploid G1 cells. **c)** The fraction of violated constraints in the 3D models of 190 haploid G1 cells at 500 kb versus 100 kb resolutions (Pearson R = 0.96). Black dots represent cells that have at least 10,000 contacts and no more than 0 violations at 1 Mb, 0.5% violations at 500 kb and 0.1% violations at 100 kb. Cells with no violations at either 500 kb or 100 kb resolution are not shown. **d)** Mean RMSD between all models of the same cell with at least 10,000 (black) and fewer than 10,000 (red) filtered contacts, for cells at 1 Mb (dots), 500 kb (open diamonds) and 100 kb (triangles) resolutions. **e)** Model reproducibility across cells. The RMSD distribution between all models for the same cell (red) compared to the RMSD distribution between models for different cells for the 126 cells with the highest quality models. The images show aligned structures of 106 ×; 1 Mb models (top), 80 × 500 kb models (centre) and 5 × 100 kb models (bottom) for NXT-1091 (38th of 126 ordered G1 cells) with a mean RMSD at the peak of the red curves. **f)** Cross-validation test. Average trans-chromosomal distance distributions, computed using a set of 200 random trans-contacts between any two chromosomes, with the contact points distributed uniformly on the two chromosomes. Red curve: trans-distances between the 5 most strongly contacting chromosome pairs in models using all supported contacts. Blue curve: same as red curve, but using the models computed with the contacts between that chromosome pair removed. Cyan curve: trans-distances of unsupported contacts in the models computed with all supported contacts present. Green curve: trans-distances of chromosome pairs in models of all cells using all supported contacts. **g)** Nucleus and chromosome shape properties in the 126 cells with high-quality models: the nuclear radius (in arbitrary units), the longest-to-shortest (a/c) semiaxis ratio of the ellipsoid fitted to the whole nucleus model, and the longest-to-shortest (a/c) and middle-to-shortest (b/c) semiaxis ratios of the ellipsoids fitted to each chromosome of the 3D models. For the nucleus size and a/c ratio, values in each model are shown, for the chromosome fitted ellipsoid semiaxis ratios the model-averaged value is shown for each cell.

**Extended Data Fig. 11:**
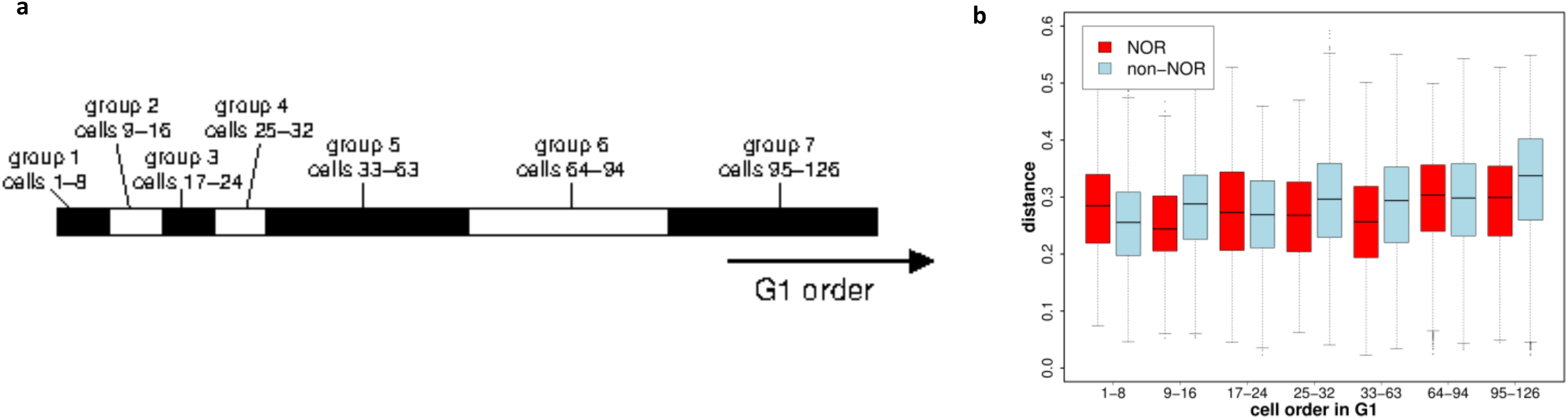
Clustering of peri-centromeric regions. **a)** The 7 cell groups in the inferred G1 time order for the 126 cells with the highest-quality models. **b)** Distances of the top third shortest distances between NOR chromosomes or non-NOR chromosomes in 7 cell groups along the inferred G1 time order. Distances are normalised by the nucleus diameter.

## References

1. Paweletz, N. Walther Flemming: pioneer of mitosis research. Nat. Rev. Mol. Cell Biol. 2, 72–75 (2001).

2. Lieberman-Aiden, E. et al. Comprehensive mapping of long-range interactions reveals folding principles of the human genome. Science 326, 289–93 (2009).

3. Sexton, T. et al. Three-dimensional folding and functional organization principles of the Drosophila genome. Cell 148, 458–72 (2012).

4. Nora, E. P. et al. Spatial partitioning of the regulatory landscape of the X-inactivation centre. Nature 5–9 (2012). doi:10.1038/nature11049

5. Dixon, J.R. et al. Topological domains in mammalian genomes identified by analysis of chromatin interactions. Nature 485, 376–80 (2012).

6. Sofueva, S. et al. Cohesin-mediated interactions organize chromosomal domain architecture. EMBO J. 32, 3119–29 (2013).

7. Rao, S. S. P. S. P. et al. A 3D Map of the Human Genome at Kilobase Resolution Reveals Principles of Chromatin Looping. Cell 159, 1665–1680 (2014).

8. Zuin, J. et al. Cohesin and CTCF differentially affect chromatin architecture and gene expression in human cells. Proc. Natl. Acad. Sci. U. S. A. 111, 996–1001 (2014).

9. de Wit, E. et al. CTCF Binding Polarity Determines Chromatin Looping. Mol. Cell 60, 676–684 (2015).

10. Flavahan, W. A. et al. Insulator dysfunction and oncogene activation in IDH mutant gliomas. Nature 529, 110–114 (2016).

11. Nagano, T. et al. Single-cell Hi-C reveals cell-to-cell variability in chromosome structure. Nature 502, 59–64 (2013).

12. Naumova, N. et al. Organization of the mitotic chromosome. Science 342, 948–53 (2013).

13. Dileep, V. et al. Topologically-associating domains and their long-range contacts are established during early G1 coincident with the establishment of the replication timing program Authors: Genome Res. 25, 1104–1113 (2015).

14. Nagano, T. et al. Single-cell Hi-C for genome-wide detection of chromatin interactions that occur simultaneously in a single cell. Nat. Protoc. 10, 1986–2003 (2015).

15. Pope, B. D. et al. Topologically associating domains are stable units of replication-timing regulation. Nature 515, 402–405 (2014).

16. Stamatoyannopoulos, J. A. et al. An encyclopedia of mouse DNA elements (Mouse ENCODE). Genome Biol. 13, 418 (2012).

17. Britton-Davidian, J., Cazaux, B. & Catalan, J. Chromosomal dynamics of nucleolar organizer regions (NORs) in the house mouse: micro-evolutionary insights. Heredity (Edinb). 108, 68–74 (2012).

18. Yaffe, E. & Tanay, A. Probabilistic modeling of Hi-C contact maps eliminates systematic biases to characterize global chromosomal architecture. Nat. Genet. 43, 1059–1065 (2011).

19. Filion, G. J. et al. Systematic protein location mapping reveals five principal chromatin types in Drosophila cells. Cell 143, 212–24 (2010).

## References

5. Dixon, J. R. et al. Topological domains in mammalian genomes identified by analysis of chromatin interactions. Nature 485, 376–80 (2012).

12. Naumova, N. et al. Organization of the mitotic chromosome. Science 342, 948–53 (2013).

14. Nagano, T. et al. Single-cell Hi-C for genome-wide detection of chromatin interactions that occur simultaneously in a single cell. Nat. Protoc. 101, 1986–2003 (2015).

20. Leeb, M. & Wutz, A. Derivation of haploid embryonic stem cells from mouse embryos. Nature 13, 1–5 (2011).

21. Leeb, M., Perry, A. C. F. & Wutz, A. Establishment and Use of Mouse Haploid ES Cells. Curr. Protoc. Mouse Biol. 5, 155–85 (2015).

22. Langmead, B. & Salzberg, S. L. Fast gapped-read alignment with Bowtie 2. Nat Methods 9, 357–l359 (2012).

23. Olivares-Chauvet, P. et al. Capturing pairwise and multi-way chromosomal conformations using chromosomal walks. Nature (2016). doi:10.1038/nature20158

24. Peric-Hupkes, D. et al. Molecular maps of the reorganization of genome-nuclear lamina interactions during differentiation. Mol. Cell 38, 603–13 (2010).

25. Wingett, S. et al. HiCUP: pipeline for mapping and processing Hi-C data. F1000Research 4, 1310 (2015).

26. Berendsen, H. J. C., van der Spoel, D. & van Drunen, R. GROMACS: A message-passing parallel molecular dynamics implementation. Comput. Phys. Commun. 91, 43–56 (1995).

27. Humphrey, W., Dalke, A. & Schulten, K. VMD: Visual molecular dynamics. J. Mol. Graph. 14, 33–38 (1996).

28. Green-Armytage, P. A Colour Alphabet and the limits of colour coding. JAIC-Journal Int. Colour… 1–23 (2010).

